# The RNA-binding protein EIF4A3 promotes axon development by direct control of the cytoskeleton

**DOI:** 10.1101/2022.03.18.484888

**Authors:** Fernando C. Alsina, Bianca M. Lupan, Lydia J. Lin, Camila M. Musso, Federica Mosti, Carly R. Newman, Lisa M. Wood, Mark Agostino, Jeffrey K. Moore, Debra L. Silver

## Abstract

The exon junction complex (EJC), nucleated by EIF4A3, is indispensable for mRNA fate and function throughout eukaryotes. Unexpectedly, we discover that EIF4A3 directly controls microtubules independent of RNA, and this is critical for neural wiring. While neuronal survival in the developing mouse cerebral cortex depends upon an intact EJC, axonal tract formation requires only *Eif4a3*. Using human cortical organoids, we demonstrate that *EIF4A3* disease mutations also impair neuronal maturation, highlighting conserved functions relevant for neurodevelopmental pathology. Employing biochemistry and molecular modeling we discover that EIF4A3 directly binds to microtubules, mutually exclusive of the RNA-binding complex. In growing neurons, EIF4A3 is essential for microtubule dynamics, and sufficient to promote microtubule polymerization and stability *in vitro*. Together, our data show that tubulin-bound EIF4A3 orchestrates microtubule dynamics, underlying key events of neuronal development. This reveals a new mechanism by which neurons re-utilize core gene expression machinery to rapidly and directly control the cytoskeleton.

**Highlights:**

- The Exon Junction Complex controls neuronal survival but only EIF4A3 directs axonal growth
- EIF4A3 controls axonal tract formation in vivo.
- Human *EIF4A3* deficient iPSC-derived cortical organoids recapitulate neuronal defects.
- EIF4A3 directly binds to microtubules to control their growth and stability in neurons.

## Introduction

The proper wiring of the mammalian cerebral cortex relies upon neuronal maturation during embryonic development (Greig et al., 2013; Lodato et al., 2015). Newborn neurons initially polarize to form axons and subsequently dendrites (Namba et al., 2015; van Beuningen and Hoogenraad, 2016). These dynamic processes depend heavily on cytoskeletal remodeling and fine-tuned gene expression, including post-transcriptional regulation (Arimura and Kaibuchi, 2007). Consistent with this, RNA-binding proteins are highly abundant in neurons and implicated in both neurodevelopmental and degenerative disorders (Hentze et al., 2018; Nussbacher et al., 2015; Schieweck et al., 2021). Yet, our understanding of how RNA-binding proteins influence neuronal development is limited. In particular, the extent to which they exclusively control RNA metabolism is largely unknown.

The RNA-binding exon junction complex (EJC) associates with 80% of the transcriptome, acting as an indispensable coordinator and guardian of mRNA fate throughout its life cycle (Le Hir et al., 2016; McMahon et al., 2016; Woodward et al., 2017). Mutations in core EJC components, *EIF4A3*, *MAGOH*, and *RBM8A*, are linked with intellectual disability and cause distinct human disorders affecting the nervous system (Albers et al., 2012; Favaro et al., 2014; Nguyen et al., 2013). For example, loss-of-function mutations in *RBM8A* and *EIF4A3* cause Thrombocytopenia-absent radius (TAR) syndrome and Richieri-Costa-Pereira Syndrome (RCPS), respectively. These divergent human disorders suggest the intriguing possibility that EJC components can also have independent functions. In the developing nervous system, EJC components are crucial for neurogenesis, where they are thought to work together (Mao et al., 2016b; Mao et al., 2015; Silver et al., 2010; Zou et al., 2015). This provides some insight into the basis for EJC-dependent disorders. However, there is a significant gap in understanding how the EJC controls brain development, as its functions in developing post-mitotic neurons are unknown (**Figure 1A**).

**Figure 1.**
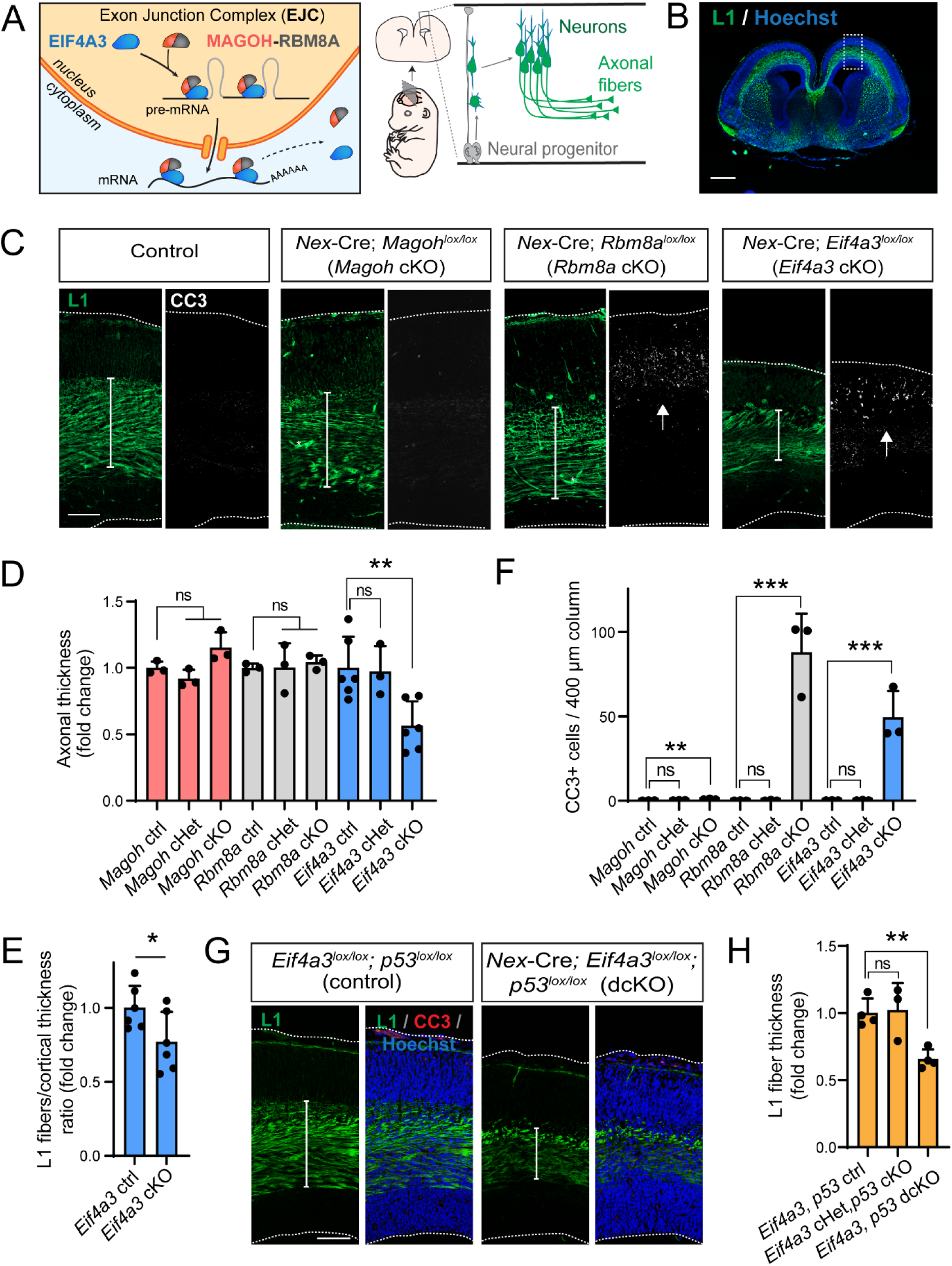
Loss of *Eif4a3* but not *Magoh* and *Rbm8a* impairs early axonal development independent of apoptosis. (**A**) Left: Cartoon depicting canonical RNA regulation by the Exon Junction Complex (EJC) composed of core components, EIF4A3 (also DDX48, blue), RBM8A (also Y14, grey), and MAGOH (red). Right: Cartoon of an embryonic mouse cortex, and question addressed in this study: How do EJC components control post-mitotic neuronal development? (**B**) Image of coronal sections from control (*Eif4a3^lox/lox^*) E17.5 mouse cortices, stained with L1-NCAM (green) and Hoechst (blue). **(C)** Higher magnification images from region indicated by dashed box in panel **(B)** stained with L1-NCAM (green) and cleaved caspase 3 (CC3, white) for indicated genotypes. White bracket, cortical axonal tract thickness. Arrow, region with apoptotic nuclei. Asterisk, background blood vessel. **(D)** Fold change of L1+ axonal thickness in E17.5 *Magoh*, *Rbm8a* and *Eif4a3* cHet and cKO cortices relative to control (*Magoh*^lox/lox^, *Rbm8a*^lox/lox^ and *Eif4a3*^lox/lox^, respectively). ns=not significant; **p=0.0065, one-way ANOVA (Dunnett’s multiple comparison test), n=3-6 embryos (2-3 litters). **(E)** Fold change of L1+ fibers/cortical thickness ratio from E17.5 *Eif4a3* cKO relative to control (*Eif4a3*^lox/lox^). **p=0.0491, one-way ANOVA (Dunnett’s multiple comparison test), n=6 embryos (3 litters). **(F)** Quantification of apoptotic cell density (CC3+) in E17.5 *Magoh*, *Rbm8a* and *Eif4a3* cHet and cKO relative to control (*Magoh*^lox/lox^, *Rbm8a*^lox/lox^ and *Eif4a3*^lox/lox^, respectively). *Magoh* ctrl vs. cKO p=0.0059; *Rbm8a* ctrl vs. cKO p=0.0004; *Eif4a3* ctrl vs. cKO p=0.001. ns=not significant; **p≤0.01; ***p≤0.001, one-way ANOVA (Dunnett’s multiple comparison test), n=3 embryos (2-3 litters). **(G)** Images of coronal sections of control (*Eif4a3*^lox/lox^, *p53*^lox/lox^) and *Eif4a3, p53* double compound cKO (dcKO) E17.5 mouse cortices, stained with L1-NCAM (green), CC3 (red) and Hoechst (blue). White bracket, cortical axonal tract thickness. **(H)** Fold change of L1+ cortical axonal thickness in E17.5 *Eif4a3*, *p53* dcHet and dcKO relative to control (*Eif4a3*^lox/lox^, *p53*^lox/lox^). **p=0.0099, one-way ANOVA (Dunnett’s multiple comparison test), n=4 embryos (4 litters). All graphs, mean + S.D. Scale bars: 500 μm (B), 100 μm (C), 50 μm (G).

In this study, we use mouse models and human cerebral organoids to dissect new requirements of EJC components in developing neurons. Remarkably, we find that while EIF4A3 is essential for axon growth *in vivo*, MAGOH and RBM8A are dispensable. Moreover, EIF4A3 is critical for formation of intracortical axonal tracts. Using biochemical and molecular modeling approaches, we discover that EIF4A3 can directly bind to microtubules and does so independent of its canonical RNA helicase functions. This interaction enables EIF4A3 to promote microtubule polymerization and stability in growing neurons. Our study demonstrates unexpected non-canonical functions for EIF4A3 in both axon formation and cytoskeletal control underlying brain wiring, which is relevant for disease.

## Results

### EIF4A3, but not MAGOH and RBM8A, is essential for early axon development

To assess how EJC components may influence neuron development, we first examined the expression of *Eif4a3*, *Rbm8a* and *Magoh* in mouse cortical neurons and neural progenitors. We used fluorescence-activated cell sorting (FACS) to purify neurons (DsRed+) and progenitors (DsRed-) from embryonic day (E) 14.5 *Dcx*::*DsRed* mouse embryos, which express DsRed under the control of a neuron-specific *Dcx* promoter (**Figure S1A**) (Wang et al., 2007). We then quantified mRNA expression of the three core EJC components, as well as *Magoh B,* a *Magoh* paralog which is 99% identical (Singh et al., 2013). Our analysis revealed that in addition to progenitors, core EJC components are also expressed in developing neurons (**Figures S1B-E**).

We next examined the requirement of the presence of each EJC component during neuronal development by conditionally removing each protein from newborn cortical excitatory neurons. For this, we crossed mice carrying floxed alleles for either *Eif4a3, Rbm8a* or *Magoh* with *Nex*-Cre, which drives Cre expression in all newborn glutamatergic neurons of the neocortex around E11.5-12.5 (Goebbels et al., 2006; Mao et al., 2016b; Mao et al., 2015; McMahon et al., 2014; McSweeney et al., 2020). Cre activity was verified by reporter expression of tdTomato in E14.5 *Nex*-Cre; *Eif4a3^lox/+^*; Ai14*^lox/+^* cortical sections (**Figure S1J**). As expected, tdTomato was only expressed in the cortical plate, where it colocalized with the neuronal marker β3-tubulin (TUBB3) but not with the progenitor marker PAX6. Next, to quantify depletion of each EJC component, we performed qPCR of FAC-sorted tdTomato+ cells from E14.5 embryonic cortices from control (*Nex*-Cre; Ai14*^lox/lox^*), *Eif4a3* (*Nex*-Cre; *Eif4a3^lox/lox^*), *Rbm8a* (*Nex*-Cre; *Rbm8a^lox/lox^*) or *Magoh* (*Nex*-Cre; *Magoh^lox/lox^*) conditional heterozygous (cHet) and conditional knockout (cKO) embryos (**Figures S1F-I**). Expression of *Eif4a3*, *Rbm8a* and *Magoh* were each significantly reduced by 50% in their respective cHets, and largely depleted in their respective cKO brains (**Figures S1F-H**). As expected, *MagohB* levels were not reduced in *Magoh* cKO (**Figure S1I**). These results demonstrate that *Nex*-Cre cKO effectively eliminates expression of EJC genes in cortical neurons.

The earliest steps of neuronal maturation involve formation of axons (van Beuningen and Hoogenraad, 2016). We thus assessed axonal organization in each cKO at E17.5, by using the pan-neuronal/axonal marker L1-NCAM (**Figure 1B**). Both *Magoh* cKO and *Rbm8a* cKO cortices showed normal axonal and cortical thickness compared to littermate controls (**Figures 1B-D** and **S1K**). In contrast, the axonal tract thickness in *Eif4a3* cKO cortices was strikingly reduced, nearly half that of control littermates (**Figures 1B-D**). As the cortex was also thinner in *Eif4a3* cKO brains (**Figure S1K**), we quantified axonal tract differences normalized to cortical thickness. This ratio was reduced, indicating that *Eif4a3* deficiency causes a disproportionate reduction in axonal fibers (**Figure 1E)**. Although *Eif4a3* cKO brains showed a striking phenotype, cHet brains were unaffected. This dosage requirement in neurons is notably different than the haploinsufficiency phenotypes of neural progenitors (Mao et al., 2016b). This suggests that *Eif4a3* may regulate neurons in a distinct fashion. Altogether, this genetic evidence demonstrates that EIF4A3 mediates early axonal development, independent of its canonical binding partners, MAGOH and RBM8A.

### Axonal growth defects precede and are independent of apoptosis in *Eif4a3* cKO mice

Defects in axon formation are frequently associated with or caused by cell death (Pemberton et al., 2021). Indeed, haploinsufficiency of core EJC components in neural progenitors induces cell death (Mao et al., 2016b). We thus analyzed apoptosis in E17.5 EJC cKO brains using the marker cleaved caspase 3 (CC3). Apoptosis was notable in the cortical plate of both *Eif4a3* and *Rbm8a* cKO mice (**Figures 1C** and **1F**). However, it was negligible in *Magoh* cKO brains, likely due to redundant expression of *MagohB* (**Figures S1E** and **S1I**). Although both *Eif4a3* and *Rbm8a* were required for neuronal viability, we only observed axonal tract defects in *Eif4a3* cKO brains (**Figure 1D).** This suggests that while the entire complex controls cell viability, only EIF4A3 is essential for axonal development.

We assessed whether *Eif4a3*-dependent axonal defects are caused by cell death. As p53 promotes apoptosis (Pemberton et al., 2021), we examined if p53 was ectopically activated in *Eif4a3* cKO brains. Indeed, relative to control, E17.5 *Eif4a3* cKO cortices showed increased p53+ nuclear staining in the cortical plate (**Figure S2A**). This suggested that neuronal death in *Eif4a3* cKO mice might be mediated by p53, thus providing a strategy to assess axonal tract development in the absence of apoptosis. We generated double conditional KO (dcKO) in which both *Eif4a3* and *p53* were deleted in excitatory neurons (*Nex*-Cre;*Eif4a3*^lox/lox^*;p53*^lox/lox^). As predicted, apoptosis was fully rescued in E17.5 *Eif4a3*; *p53* dcKO brains (**Figure S2B).** Notably, relative to littermate controls, L1+ axonal tract thickness was still significantly reduced in *Eif4a3; p53* dcKO, to a comparable extent to *Eif4a3* cKO alone (**Figures 1G, 1H** and **S1L**). These data strongly indicate that apoptosis does not cause the axonal tract defect in E17.5 *Eif4a3* cKO brains.

As neurons are born, they begin to polarize and generate axons (Namba et al., 2015). To assess if *Eif4a3* is required for these early stages of axonal development, we assessed E14.5 brains. Single molecule FISH in E14.5 *Eif4a3* cKO brains confirmed reduced *Eif4a3* expression in the cortical plate (**Figure S2C)**. First, we assessed if apoptosis was evident at this earlier stage. E14.5 *Eif4a3* cKO brains showed virtually no apoptosis, especially as compared to E17.5, with just 2 apoptotic cells versus >50, respectively (compare **Figure S2D** to **Figure 1F**). However, both E14.5 *Eif4a3* cKO and *Eif4a3; p53* dcKO brains displayed substantially disorganized axonal tracts compared to control, evidenced by shorter (<20 μm) and less defined axons (**Figures S2E-G**). In contrast, axonal length distribution was normal in *Rbm8a* cKO and *Magoh* cKO E14.5 brains (**Figures S2H** and **S2I**). Altogether, analyses at two developmental stages indicate that *Eif4a3* has distinct genetic requirements from *Rbm8a* and *Magoh* in axonal development which precede and are independent of apoptosis.

### EIF4A3 is autonomously required for neuronal polarization and axonal growth, independent of RBM8A and MAGOH

Newborn neurons first polarize to form axons (van Beuningen and Hoogenraad, 2016). Given defects in axonal tract integrity *in vivo*, we predicted that *Eif4a3* may be required for neuronal polarization and axonal growth. We thus measured axon length in primary neuronal cultures generated from E15.5 control and *Eif4a3* cKO cortices (**Figures 2A-C**). EIF4A3 protein expression was reduced in day *in vitro* 2 (DIV2) cKO cultures (**Figure S3A)**. Strikingly, *Eif4a3* cKO showed a 32% average reduction in axonal length compared to control (**Figure 2C**). Moreover, the proportion of polarized cKO neurons (displaying a clear longest axon projection) was reduced by 38% (**Figure S3B**). Importantly, these developing cKO neurons were not apoptotic (**Figure S3C**). These results demonstrate a cell-autonomous requirement of *Eif4a3* in axonal formation and reinforce the conclusion that axonal impairments precede apoptosis.

**Figure 2.**
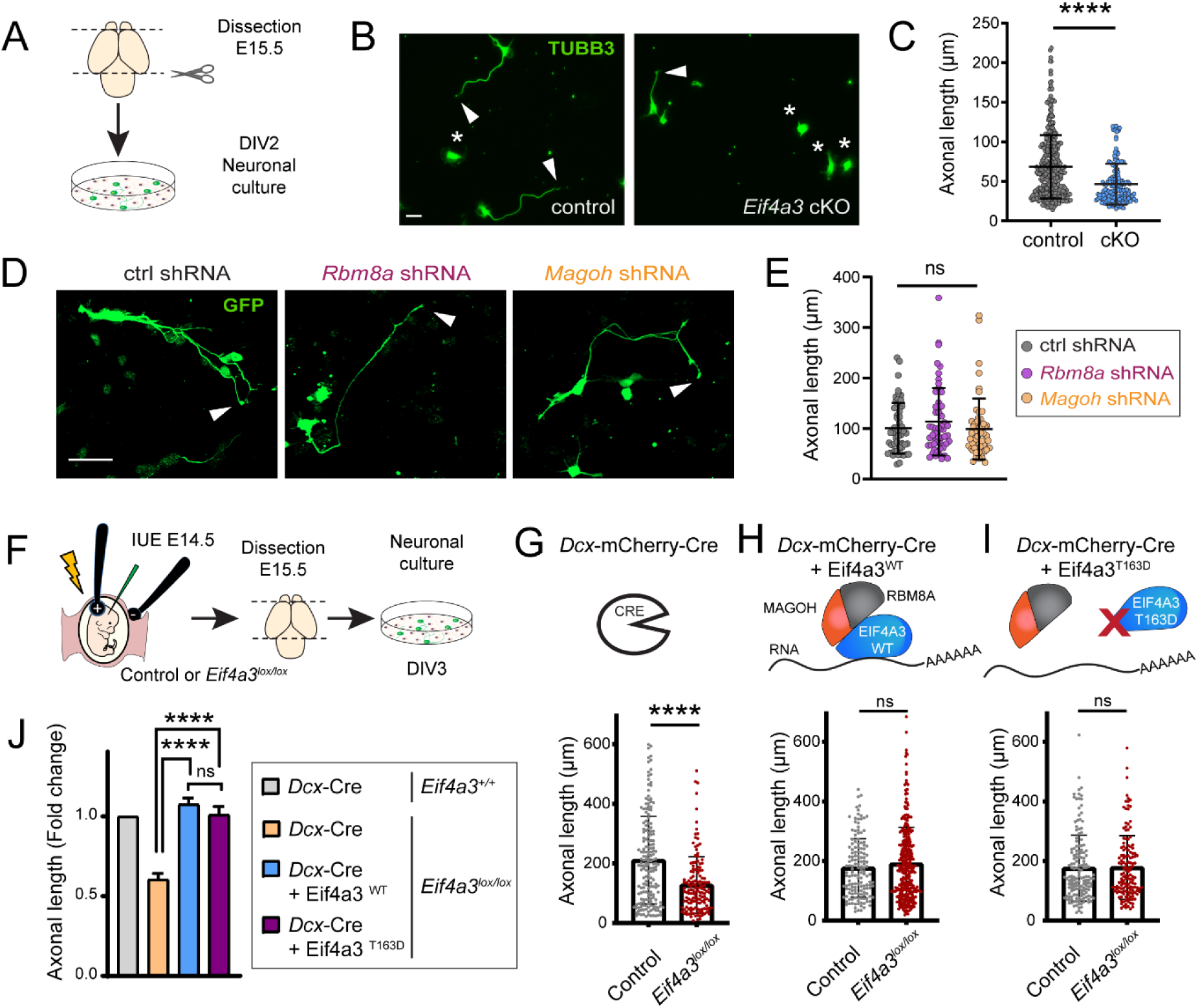
EIF4A3-mediated axonal development is EJC and RNA independent. **(A)** Cartoon depicting E15.5 cortical neuronal culture preparation from mouse embryos, fixed at DIV2. **(B)** Images of neuronal cultures stained against TUBB3 (β3-tubulin). Arrowheads, axons of polarized neurons. Asterisks, unpolarized neurons. **(C)** Quantification of axonal length of low-confluency neuronal cultures from control and *Eif4a3* cKO littermates. ****p<0.0001, unpaired t-test (two tailed), n=377 control neurons (4 embryos; 3 litters) and n=151 cKO neurons (3 embryos;3 litters). **(D)** Images of *wild type* neurons from high-confluency E15.5 cultures, co-electroporated with CAG-GFP and scrambled shRNA, *Rbm8a* shRNA or *Magoh* shRNA at DIV0 and fixed at DIV2. Arrows, axons. **(E)** Quantification of axonal length of neuronal cultures; ns=not significant, one-way ANOVA (Dunnett’s multiple comparison test), n=55 neurons for control shRNA and *Rbm8a* shRNA, n=53 for *Magoh* shRNA (3 experiments each). **(F)** Schematic of rescue experiments in which E14.5 control (*Eif4a3*^+/+^) or *Eif4a3*^lox/lox^ littermate embryos were *in utero* electroporated with CAG-GFP, *Dcx*-mCherry-Cre, and constructs in (**G-I**). **(G-I)** Axonal length quantification from GFP+, mCHERRY+, FLAG+ DIV3 neuronal cultures of control (*Eif4a3*^+/+^ or *Eif4a3*^lox/+^) or *Eif4a3*^lox/lox^ littermate embryos *in utero* electroporated with CAG-GFP and *Dcx*-mCherry-Cre (G) CAG-GFP, *Dcx*-mCherry-Cre and CAG-3x-flag- EIF4A3^WT^ (H), and CAG-GFP, *Dcx*-mCherry-Cre and CAG-3x-flag- EIF4A3^T163D^ (I).****p<0.0001, ns=not significant, unpaired t-test (two tailed), n=143-310 neurons (2-5 embryos; 2-3 litters). **j,** Fold change of axonal length relative to control (*Eif4a3*^+/+^) from *Eif4a3*^lox/lox^ embryos *in utero* electroporated with indicated constructs (data from **G,H,I**). ****p<0.0001, ns=not significant, one-way ANOVA (Tukey’s multiple comparison test), n=143-310 neurons (2-5 embryos; 2-3 litters). All graphs, mean + S.D. Scale bars, 20 μm (B), 50 μm (D).

We next assessed whether axon growth and polarization defects are a general consequence of impaired EJC function. To address this, wild type E15.5 neurons were transfected at DIV0 with shRNAs against *Rbm8a* and *Magoh* and cultured for two days. Both shRNAs significantly depleted their respective targets (**Figures S3D** and **S3E).** However, DIV2 axonal growth was unaffected when either gene was downregulated (**Figures 2D** and **2E**). This further supports the unique requirement of *Eif4a3* for axonal development, and indicates it functions in an EJC-independent fashion.

The cellular functions of EIF4A3 are typically attributed to its ability to bind both RNA and proteins of the EJC. To further assess the extent to which these interactions are required for EIF4A3-mediated neuronal maturation, we investigated whether expression of EIF4A3^WT^ or a previously described point mutant, EIF4A3^T163D^, which fails to bind the EJC and RNA (Ryu et al., 2019), would restore axon formation in *Eif4a3* mutant neurons. We confirmed that EIF4A3^T163D^ fails to bind MAGOH (**Figure S3F**). E14.5 *Eif4a3^+/+^* (control) or *Eif4a3^lox/lox^* littermate embryos were *in utero* electroporated (IUEd) with one of the following: (1) *Dcx*-Cre-mCherry; (2) *Dcx*-Cre-mCherry and flag-EIF4A3^WT^ (positive control); or (3) *Dcx*-Cre-mCherry and flag-EIF4A3^T163D^ (**Figures 2F-I**). At E15.5, primary neuronal cultures were generated and subsequently fixed at DIV3 (**Figure 2F**). Importantly, neither EIF4A3^WT^ or EIF4A3^T163D^ expression caused an over- expression phenotype in *wild-type* neurons, as evidenced by no substantial change in axonal and neurite outgrowth (**Figures S3G** and **S3H**). Further, both tagged proteins showed similar localization to the cell body and neuronal projections (**Figures S3I** and **S3J**). Axonal length was reduced in *Eif4a3^lox/lox^* neurons expressing *Dcx*-mCherry-Cre, and to a similar extent as *Eif4a3* cKO neurons (**Figures 2G** and **2C**). As expected, expression of EIF4A3^WT^ rescued these axonal length defects of *Eif4a3* deficient neurons (**Figure 2H**). We then assessed whether EIF4A3^T163D^ would rescue these defects. Indeed, both the axonal defect, as well as neurite outgrowth were rescued to a comparable degree when EIF4A3^T163D^ was expressed in *Eif4a3*-deficient neurons (**Figures 2I, 2J,** and **S3K-M**). Taken together, these data demonstrate that EIF4A3 controls early axonal and neurite outgrowth and does so in an EJC- and RNA- independent manner.

### Human cortical organoids modeling *EIF4A3*-associated disease exhibit abnormal neuronal growth and maturation

Having found that *Eif4a3* loss-of-function disrupts early axon development, we next asked whether this requirement is conserved in humans and relevant for disease. Similar to mice, human *EIF4A3* is expressed in progenitors and neurons of the developing cortex (Nowakowski et al., 2017). Notably, *EIF4A3* noncoding mutations in its 5′ untranslated region (UTR) cause RCPS, a craniofacial disorder in which patients also exhibit language and learning impairments, and mild microcephaly. Unaffected persons typically have 6-9 repeats of 18-20 nucleotide-long motifs within the *EIF4A3* 5ʹUTR (Hsia et al., 2018), whereas most affected individuals have 14-16 expanded repeats, resulting in hypomorphic *EIF4A3* expression (Favaro et al., 2014) (**Figures S4A** and **S4B**). While the craniofacial basis of RCPS has been investigated (Miller et al., 2017), the neurodevelopmental etiology for RCPS brain deficits is unknown.

We thus used control and RCPS patient induced pluripotent stem cell (iPSC) lines to generate cortical organoids (Yoon et al., 2019) (**Figures 3A** and **S4A)**. At day 35 (D35) of differentiation, both control and patient-derived organoids had neural rosette structures bearing β3-tubulin+ neurons and Sox2+ neural progenitors (**Figure S4C**). As further characterization, in both control and RCPS D25 organoids, we observed downregulation of pluripotency markers *OCT4* and *NANOG,* absence of mesoderm *BRACH* and endoderm *SOX17* markers, and upregulation of forebrain transcripts *FOXG1* and *PAX6* (**Figures S4D-I**). Additionally, we generated an isogenic control line from one of the RCPS iPSCs (**Figure S4J**). Importantly, we quantified a ∼50% reduction in *EIF4A3* mRNA from RCPS patient-derived cortical organoids compared to controls, similar to that previously seen in RCPS patient-derived neural crest cells (Miller et al., 2017) (**Figure 3B**). These data demonstrate that control and patient iPSC lines can be reliably differentiated into cortical organoids to model aspects of early human brain development.

**Figure 3.**
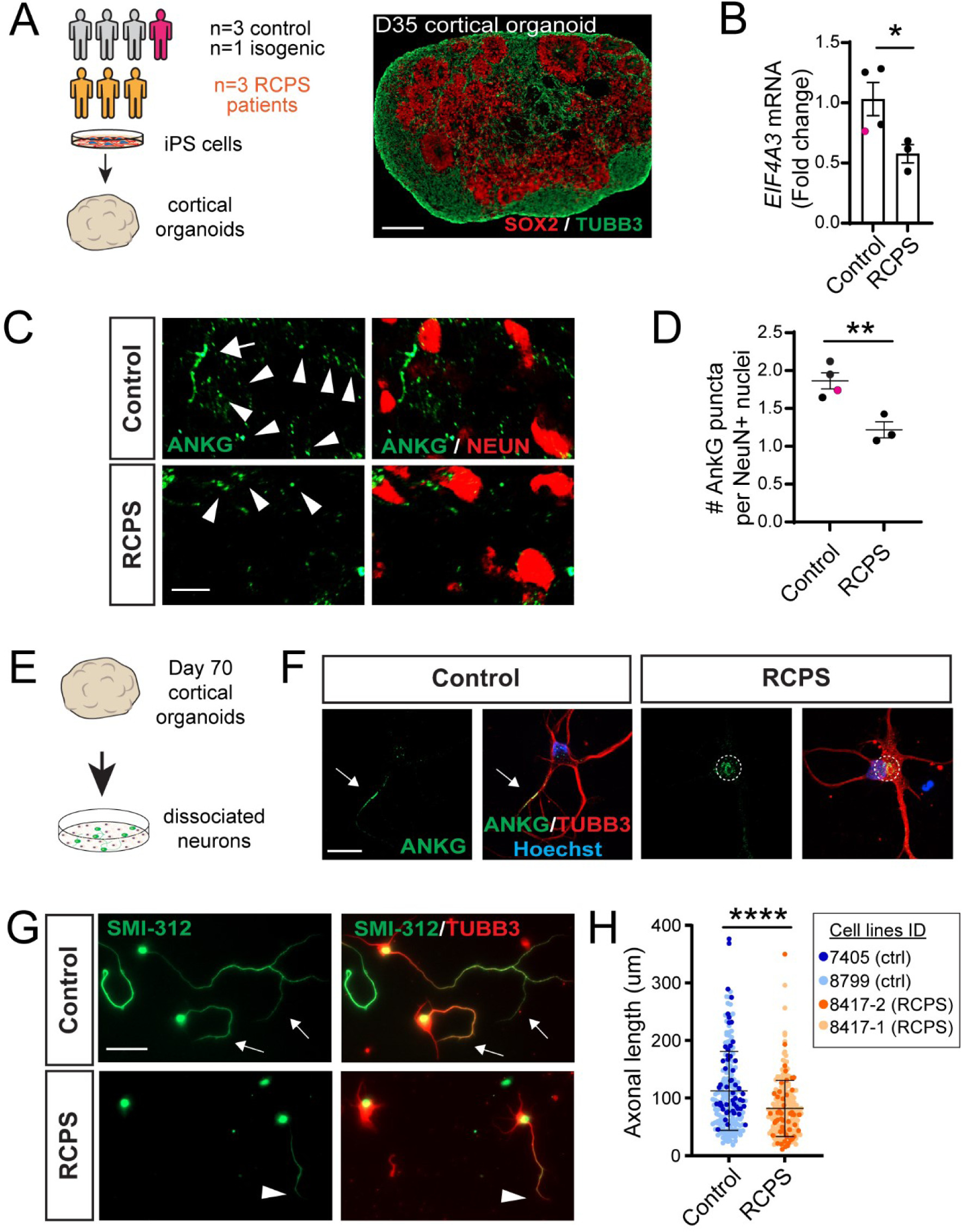
Impaired axonal development in human cortical organoids derived from RCPS patients. **(A)** Left: Cartoon of the generation of human brain cortical organoids from 3 control, 1 isogenic, and 3 Richieri-Costa-Pereira Syndrome (RCPS) patient-derived iPSCs. Right: Image of a day 35 (D35) control cortical organoid stained against SOX2 (red) and TUBB3 (green). **(B)** RT-qPCR analysis of *EIF4A3* mRNA levels in control and RCPS organoids. Magenta, isogenic control. All organoids D25 except RCPS line F6099-1 (D49). *p=0.0487, unpaired t-test (two tailed), n=4 cell lines for control (including isogenic); n=3 cell lines for RCPS (3-9 organoids;2 differentiations). **(C)** Images from cryosections of D35 cortical organoids, stained for Axon Initial Segment (AIS) marker ANKYRIN-G (ANKG, green) and nuclear neuronal marker NEUN (red). Arrows, AIS in longitudinal plane. Arrowheads, AIS puncta. **(D)** Quantification of ANKG puncta per NEUN+ nuclei in control and RCPS patient derived organoids. Magenta, isogenic control. **p=0.0087, unpaired t-test (two tailed), n=4 control (including isogenic); n=3 RCPS (8-10 organoids each; 2 differentiations). **(E)** Schematic of cortical organoid dissociation to generate 2D neuronal cultures. **(F)** Images of DIV7 neuronal cultures from D71-73 control and RCPS cortical organoids, stained with ANKG (green), TUBB3 (red) and Hoechst (blue). Arrow, defined AIS. White dashed circle, diffuse somal ANKG staining. **(G)** Images of DIV4 neuron cultures from D71- 73 control and RCPS cortical organoids, stained with the axonal marker SMI-312 (green) and TUBB3 (red). Arrows, strong SMI-312 signal in distal part of the axon. Arrowhead, shorter axon with dimmer SMI-312 staining. (**H**) Quantification of axonal length across 2 controls and 2 RCPS lines. n=203 control neurons (n=2 cell lines) and n=222 cKO neurons (n=2 cell lines). p<0.0001, unpaired t-test. All graphs, mean + S.D. Scale bars, 200 μm (A), 10 μm (C), 20 μm (F), 50 μm (G).

We next evaluated if RCPS organoids exhibit early axon development defects. In order to discretely label neurons, we stained D35 organoids for the Axon Initial Segment (AIS) marker AnkyrinG (AnkG). The AIS is a hallmark of early neuronal maturation, after axon formation (Leterrier and Dargent, 2014). Compared to control, AnkG puncta were significantly reduced in RCPS organoids (**Figures 3C** and **3D**). Notably, this AnkG staining was also reduced in *Eif4a3* cKO brains (**Figure S4K**), indicating a conserved defect in neuronal maturation. To further resolve AnkG localization in individual human neurons, we dissociated control and RCPS organoids and generated neuronal cultures (**Figure 3E**). While control neurons successfully began to form the AIS in DIV7 cultures, RCPS neurons showed diffuse and somatic AnkG staining (**Figures 3F** and **S4L**). Moreover, growth of RCPS neurons was impaired, as evidenced by both shorter axons and reduced axonal expression of the pan-axonal neurofilament marker SMI-312 (**Figure 3G**). We quantified shorter axonal length in RCPS neurons compared to control (**Figure 3H**). These results indicate that RCPS human organoids show defective neuronal development, a phenotype which may be relevant for human disease. Taken together with our findings using cKO mice, this demonstrates that *EIF4A3* has conserved requirements for neuronal growth and maturation.

### EIF4A3 is required for *in vivo* axon formation and integrity of axonal tracts

Our results show that *Eif4a3* is essential for early axonal development, raising the question of whether it impacts cortical wiring. To evaluate this, we focused specifically on commissural axonal tracts. While the earliest-born projection neurons project sub-cortically, callosal projection neurons born later extend axons to the contralateral hemisphere (Greig et al., 2013; Lindwall et al., 2007). Additional axonal tracts connecting the two hemispheres include the anterior commissures and the hippocampal commissures.

We thus performed an anatomical analysis of whole brains using an adapted CUBIC tissue clearing protocol coupled with Light Sheet Microscopy (Susaki et al., 2015) (**Figure 4A**). We evaluated E14.5 heads from *Eif4a3* cHet (control) and cKO mice carrying *Ai14^lox/+^*, in which tdTomato labels glutamatergic neurons including their axonal projections. We then analyzed the anterior commissure, the earliest and main intracortical axonal tract to form in the embryonic mouse brain at this stage (Lindwall et al., 2007). *Eif4a3* cKO brains showed defective projections to the midline, with a concomitant 55% reduction in commissure volume (**Figures 4B** and **4C, Video S1).** To comprehensively evaluate the three main commissural axonal tracts, we then analyzed brains at E17.5, when all connections have started to form (**Figures S5A-C**). For the corpus callosum, hippocampal commissures, and anterior commissures, we observed a substantial volumetric reduction (**Figures 4D-F**, and **S5D; Video S2**). Taken together with L1 staining, these findings demonstrate that *Eif4a3* is crucial for integrity of commissural axonal tracts in the neocortex.

**Figure 4.**
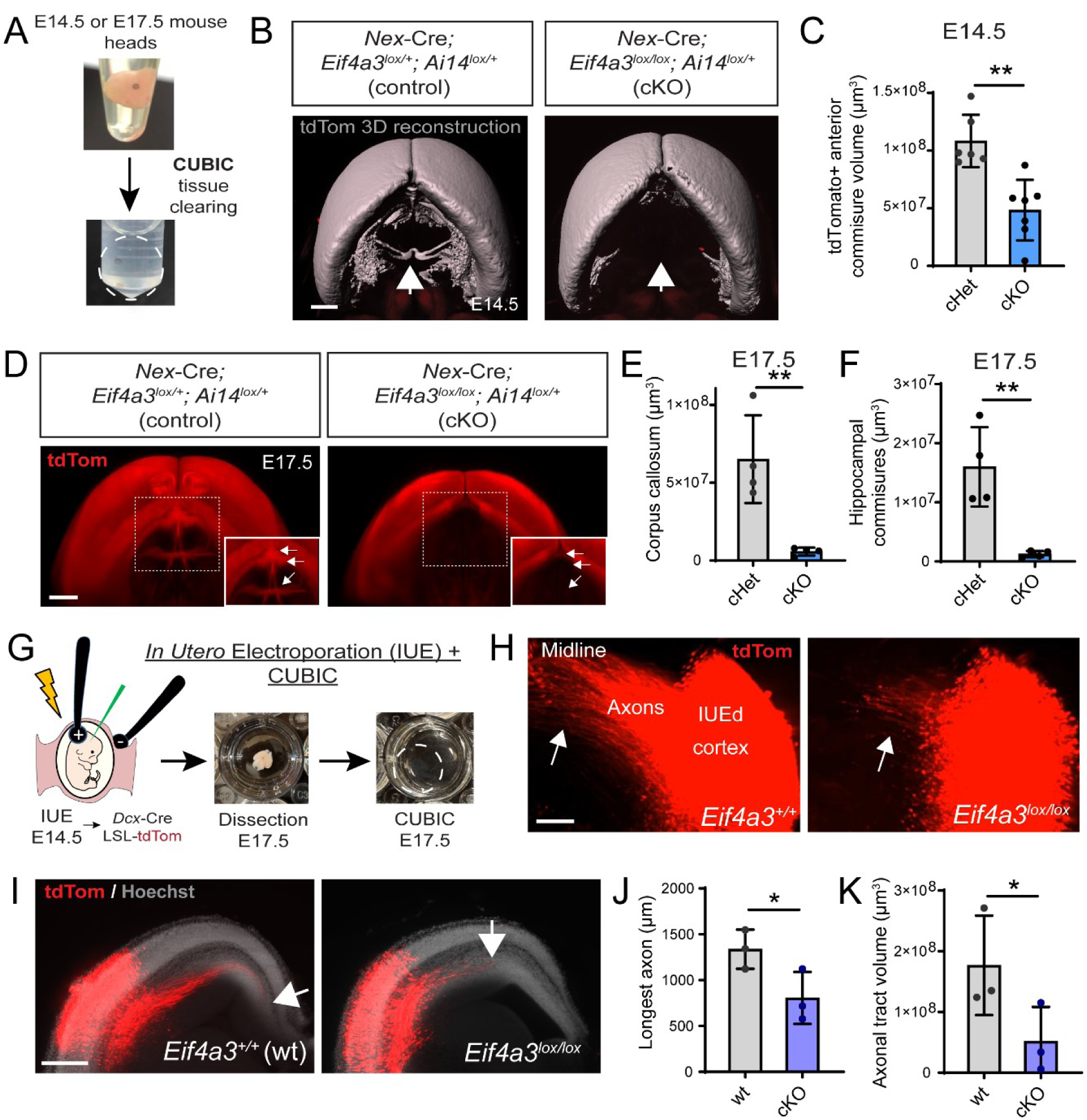
*Eif4a3* is required for axonal tract formation *in vivo*. **(A)** Representation of CUBIC tissue clearing, with dashed circle depicting the cleared embryonic mouse head. **(B)** 3D reconstructions of tdTomato+ E14.5 mouse brains from *Eif4a3* cHet (control) and *Eif4a3* cKO (caudal view). Grey color, tdTomato signal used to generate volumetric quantifications. Arrowheads, anterior commissures. **(C)** Anterior commissures volume in E14.5 *Eif4a3* cHet (control) and *Eif4a3* cKO brains. **p=0.0011, unpaired t-test (two tailed), n=6 control brains (4 litters) and n=7 *Eif4a3* cKO brains (3 litters). **(D)** 3D reconstructions of tdTomato+ E17.5 mouse brains from *Eif4a3* cHet (control) and *Eif4a3* cKO (caudal view). Box (white dash) depicts location of corpus callosum, hippocampal commissures, and anterior commissures (indicated by arrows from top to bottom, respectively). **(E,F)** Corpus callosum (**E**), and Hippocampal commissures (**F)**, volume in E17.5 *Eif4a3* cHet (control) and *Eif4a3* cKO brains. **p=0.0057 (**E**), **p=0.0047 (**F**), unpaired t-test (two tailed), n=4 control and *Eif4a3* cKO brains (from 2 litters each). **(G)** Schematic of experimental design to sparsely label neurons and their projections in 3D, including *in utero* electroporation (IUE) at E14.5, brain dissection and clearing by CUBIC at E17.5 (dashed circle). **(H)** 3D reconstructed cortices from *Eif4a3*^+/+^ (control) and *Eif4a3*^lox/lox^ embryos electroporated with *Dcx*-Cre-GFP and lox-stop-lox-tdTomato. Arrows, groups of axons extending to the midline after 3 days of electroporation. **(I)** Optical planes of 3D reconstructed brains from *Eif4a3*^+/+^ (control) and *Eif4a3*^lox/lox^ embryos labeled as in (**H**). Arrows, the longest groups of axons extending to the midline after 3 days of electroporation. **(J)** Axonal length of the average of five electroporated tdTomato+ longest axons measured from the cortical plate to the midline in *Eif4a3*^+/+^ (control) and *Eif4a3*^lox/lox^ embryos. *p=0.0302, unpaired t-test (one tailed), n=3 control and *Eif4a3* cKO brains (from 2 litters). **(K)** tdTomato+ axonal fiber volume measured from the electroporated region in the cortical plate to the midline in *Eif4a3*^+/+^ (control) and *Eif4a3*^lox/lox^ embryos. *p=0.0476, unpaired t-test (one tailed), n=3 control and Eif4a3 cKO brains (2 litters). All graphs, mean + S.D. Scale bars: 500 μm (B,I). 700 μm (D), 250 μm (H).

To better understand the temporal requirements of *Eif4a3* for axon tract formation, we targeted *Eif4a3* expression in a sparse, acute manner by using *in utero* electroporation (IUE) coupled with CUBIC (**Figure 4G**). E14.5 *Eif4a3*^+/+^ (control) or *Eif4a3^lox/lox^* brains were electroporated with *Dcx*-Cre-GFP and a Cre reporter (LSL-tdTomato) to target and label *Eif4a3* expression specifically in newborn callosal neurons. E17.5 brains were then imaged to visualize callosal neurons, whose axons project to the midline to form the corpus callosum (**Figures 4H**, and **4I; Video S3).** Compared to control, *Eif4a3^lox/lox^* electroporated brains had a significant 40% reduction in the average axonal length extending towards the midline and a 71% decrease in the total volume of axonal tracts (**Figures 4J** and **4K**). Taken together, these findings indicate that *Eif4a3* is specifically required in excitatory neurons for *in vivo* axonal formation and the development of brain commissures.

### EIF4A3 is essential for microtubule growth and stability in neurons

Our findings thus far demonstrate that EIF4A3 underlies axonal tract formation and does so independent of the EJC. In order to understand how EIF4A3 may uniquely control axonal development, we next evaluated its sub-cellular localization in developing neurons. For this, we used DIV2 primary neurons from E15.5 mouse cortices, three different antibodies, and two super-resolution microscopes (SIM, STED). EIF4A3 not only localized to soma but was also highly enriched in neuronal projections (β3-tubulin/Tubb3 positive) (**Figures 5A, 5B,** and **S6A**). In comparison, MAGOH and RBM8A were primarily nuclear/somatic, showing a lower axon:nuclear ratio (**Figures 5A** and **5B**). This is consistent with previously described localization of the heterodimer in other cell types (Chan et al., 2004; Kataoka et al., 2000; Le Hir et al., 2001). The enriched localization of EIF4A3 in neurites is consistent with biochemical studies of the EJC; while nuclear EIF4A3 is primarily EJC-bound, cytoplasmic EIFA3 is not (Chan et al., 2004; Singh et al., 2012). Further, knockdown studies demonstrate that EIF4A3 is stable outside of the EJC and its stable association with RNA requires the MAGOH-RBM8A heterodimer (Andersen et al., 2006; Bono et al., 2006; Gehring et al., 2009; Wang et al., 2014). Taken together with our genetics and rescue experiments, this suggests that there is a pool of non-EJC and non-RNA bound EIF4A3 in neuronal projections which controls development via non-canonical mechanisms.

**Figure 5.**
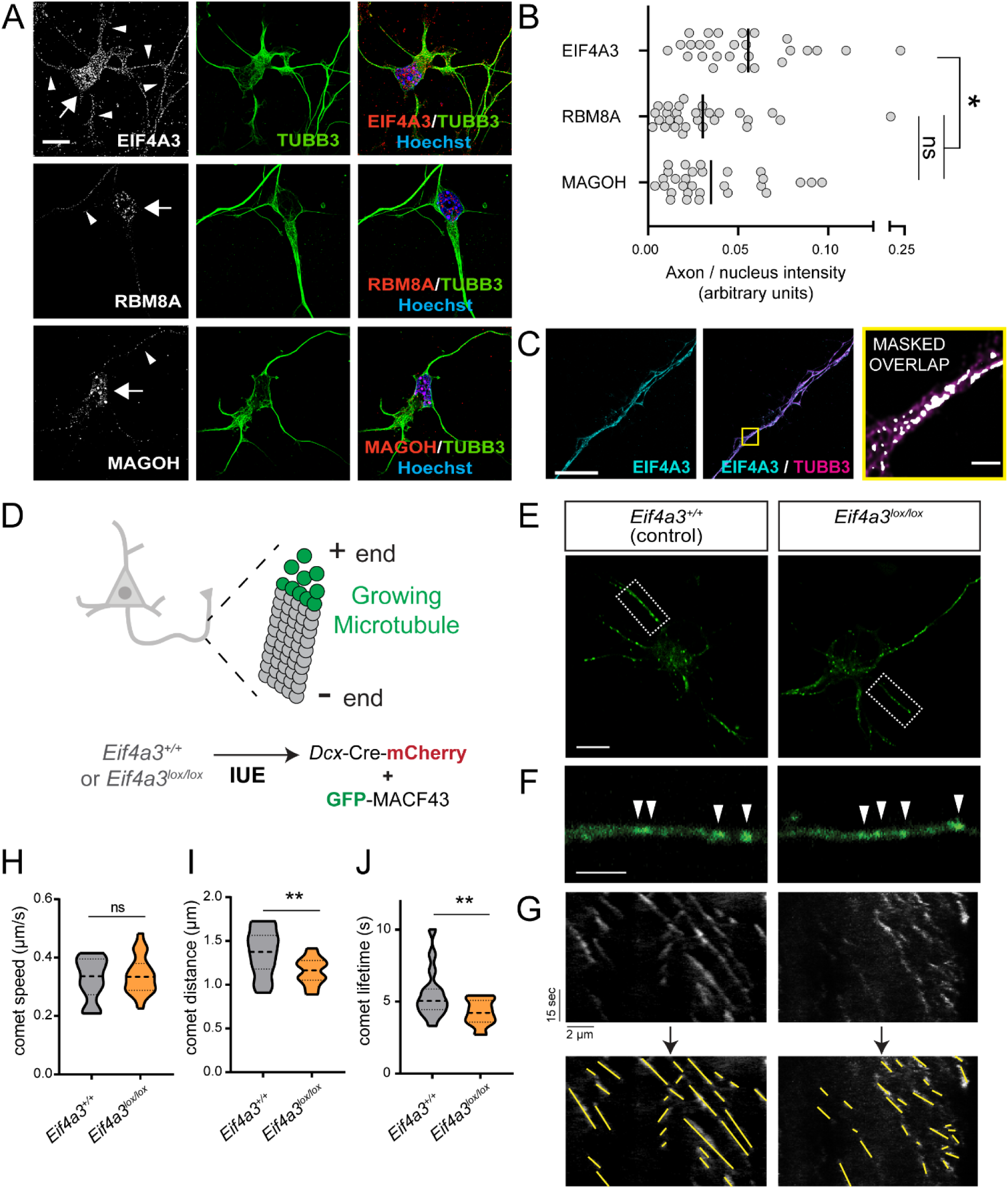
EIF4A3 is enriched in neuronal projections and required for microtubule growth and stability in developing neurons. **(A)** SIM super-resolution images of DIV2 cortical neurons from wild type E15.5 neuronal cultures, stained for either EIF4A3, RBM8A or MAGOH (red) and TUBB3 (green) and Hoechst (blue). Arrows, nucleus; arrowheads, neuronal projections. **(B)** Quantification of axonal/nuclear signal for each indicated protein. *p=0.0122 (EIF4A3 vs. RBM8A) and p=0.0463 (EIF4A3 vs. MAGOH), ns=not significant, one-way ANOVA (Tukey’s multiple comparison test), n=27 neurons for EIF4A3 staining, n=28 neurons for RBM8A staining, n=29 neurons for MAGOH staining. **(C)** STED super-resolution images of a DIV2 cortical axon from wild type E15.5 neuronal cultures, stained for EIF4A3 (cyan) and TUBB3 (magenta). Yellow box, higher magnification of middle panel; white represents a mask overlap of EIF4A3 with TUBB3. **(D)** Schematic of microtubule live imaging of DIV2 neurons prepared from E14.5 control (*Eif4a3*^+/+^) or *Eif4a3*^lox/lox^ littermate embryos *in utero* electroporated with GFP-MACF43 and *Dcx*-mCherry- Cre. **(E)** Representative snapshots of live imaging of DIV2 cortical neurons from control (*Eif4a3*^+/+^) or *Eif4a3*^lox/lox^. GFP puncta indicate GFP-MACF43+ growing microtubules. **(F)** Higher magnification image from region in panel **E**. Arrowheads, growing microtubule comets visualized as GFP+ puncta. **(G)** Representative kymograph reconstructions from live imaging of comets (growing microtubules) in neurites in DIV2 neuronal cultures. Lower images, tracings of individual comet events. **(H-J)** Quantification of comet speed (**H),** comet distance **(I),** comet lifetime (**J)** from neuronal cultures from control (*Eif4a3*^+/+^) or *Eif4a3*^lox/lox^ littermates. **p=0.0042 (**I**), **p=0.0077 (**J**), unpaired t-test (two tailed), n=15 control electroporated neurons (1307 traced comets from 2-6 neurites per neuron; 3 embryos; 2 litters) and n=20 *Eif4a3* lox/lox electroporated neurons (2049 traced comets from 2-6 neurites per neuron; 4 embryos; 3 litters). Graph on **(B)**, mean + S.D; graphs on (**H-J**) truncated violin plots with median (thicker dashed-line) ± quartiles (thinner dashed-lines). All graphs, mean + S.D. Scale bars: 10 μm (A, C-left, E), 1 μm (C-right); 2.5 μm (F).

Having established that EIF4A3 controls axon development independent of the EJC, we sought to understand the molecular mechanisms at play. Neuronal projections are highly enriched for cytoskeletal components, and both neuronal polarization and axon formation rely heavily on microtubule dynamics (van Beuningen and Hoogenraad, 2016). Notably, super-resolution microscopy showed EIF4A3 and β3-tubulin colocalized in neuronal axons and neurites at nanometer resolution (**Figure 5C**). This suggests that EIF4A3 may influence microtubules in neurites.

As microtubule dynamics are integral to proper axon formation, we monitored growing microtubules in *Eif4a3* deficient neurons. For this, we used GFP-tagged MACF43, a fragment of the MACF2 protein that binds to EB proteins at the plus ends of growing microtubules (Honnappa et al., 2009) (**Figure 5D**). GFP-MACF43 and *Dcx*-Cre-mCherry constructs were electroporated in E14.5 *Eif4a3^+/+^* (control) or *Eif4a3^lox/lox^* brains. E15.5 primary neuronal cultures were generated, and we performed live cell imaging of GFP+ comets in DIV2 neurons, a stage when neurons extend multiple immature neurites to achieve axon specification (**Figure 5E** and **5F**). We tracked thousands of comets from multiple neuronal projections and neurons, and generated kymographs to analyze their behavior (**Figure 5G**). Neither the total number of comets in the soma and neurites nor comet speed or microtubule growth orientation were affected by *Eif4a3* loss (**Figures 5H** and **S6B-D**). However, both comet distance and lifetime were significantly reduced following *Eif4a3* deletion (**Figures 5I** and **5J; Video S4**). This indicates that EIF4A3 is essential for microtubule growth and stability.

These microtubule defects could precede or coincide with altered axonal growth. To discriminate between these possibilities, we quantified axonal growth at DIV2. This is one day earlier than previously identified axonal defects (**Figures 2G** and **S3K**). Importantly, at this stage, both axonal and neurite outgrowth were normal in *Eif4a3* deficient neurons (*Eif4a3^+/+^* and *Eif4a3^lox/lox^* neurons electroporated with *Dcx*-Cre-mCherry) (**Figures S6E** and **S6F**). Taken together with microtubule imaging, these experiments indicate that aberrant microtubule dynamics clearly precede neuronal growth defects. We then used Taxol stabilization of microtubules in cKO neurons, which partially recovered axon length (**Figure S6G**). This is consistent with the possibility that microtubule instability underlies aberrant axon formation. Altogether, these results strongly suggest that EIF4A3 is required for microtubule stability to promote axonal and neurite outgrowth.

### EIF4A3 directly binds to microtubules independent of RNA and mutually exclusive of the EJC

How does EIF4A3 loss impair microtubule dynamics? As EIF4A3 controls microtubules stability and colocalizes with β3-tubulin in neurites, we reasoned that EIF4A3 may biochemically associate with tubulin. To assess this possibility, we performed *in vivo* co-immunoprecipitation of the neuronal specific subunit β3-tubulin (TUBB3) and EIF4A3 using E16.5 mouse neocortices (**Figure 6A**). We also included co-immunoprecipitations of MAGOH and RBM8A, for comparison, since neither showed significant neurite localization. EIF4A3, but not MAGOH and RBM8A, was specifically precipitated by TUBB3 (**Figure 6B**). Reverse co-immunoprecipitation using EIF4A3 antibodies showed a similar association with TUBB3 (**Figure 6C)**. These data indicate that EIF4A3, but not MAGOH or RBM8A, associates *in vivo* with tubulin in developing neurons. As these experiments were performed in clarified lysates under cold conditions where tubulins are not polymerized, our data further suggest that EIF4A3 may be competent to bind free tubulin heterodimers.

**Figure 6.**
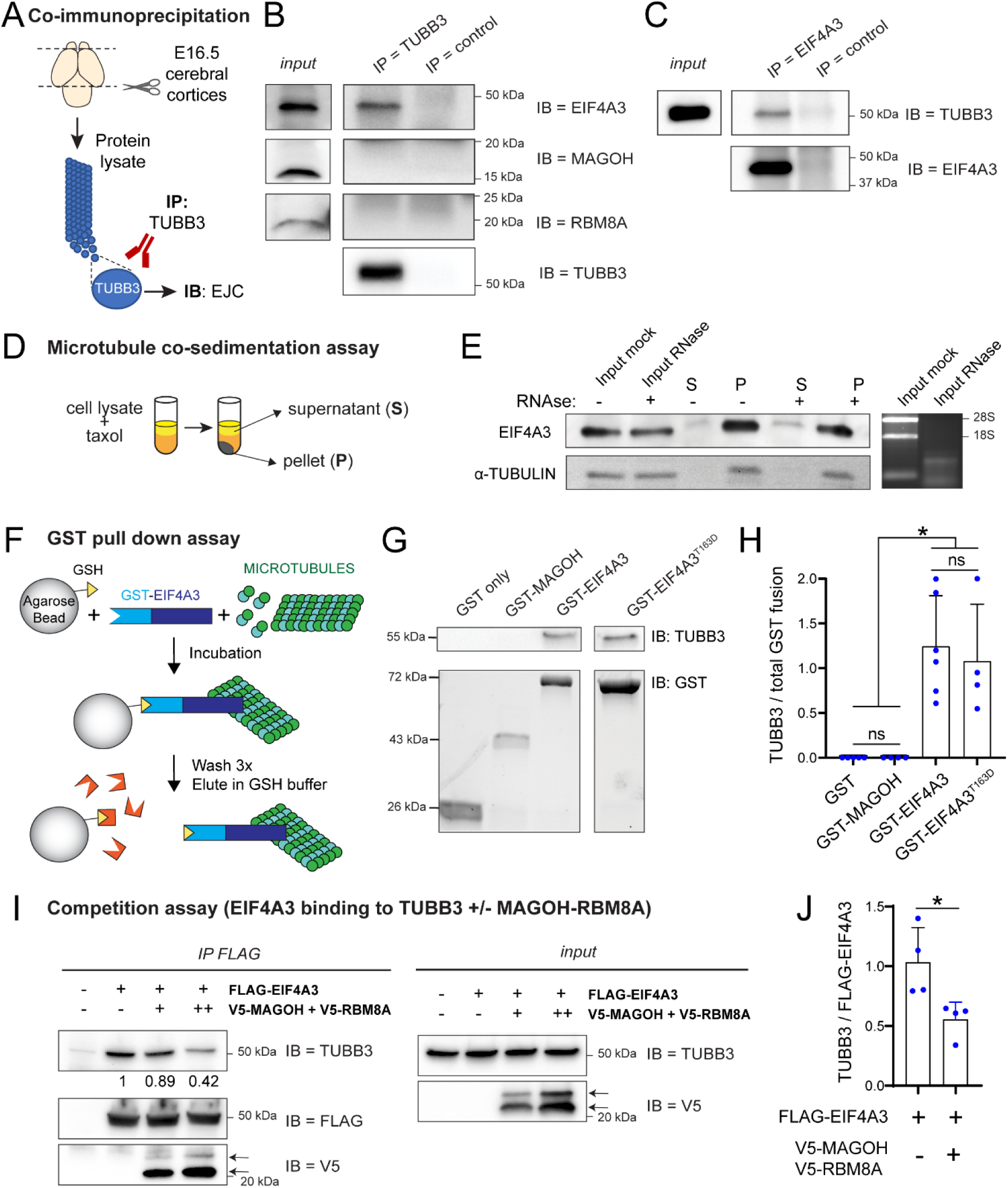
EIF4A3 directly binds to microtubules in a mutually exclusive and EJC- independent manner. **(A)** Cartoon of co-immunoprecipitation (co-IP) from E16.5 mouse cortices. **(B)** Representative immunoblots (IB) depicting co-IP of TUBB3 or FLAG (control) with core EJC components (EIF4A3, MAGOH, and RBM8A), re-probing of the same blot with TUBB3 antibody, and input lysates. n=2 experiments. **(C)** Representative co-IPs of EIF4A3 or RFP (control) with TUBB3, re-probing of the same blot with EIF4A3 antibody, and input lysates. n=4 experiments. **(D)** Cartoon of a microtubule co-sedimentation assay showing the supernatant (S, yellow), glycerol cushion (blue) and microtubule pellet (P, tan). **(E)** Representative immunoblot depicting mock and RNase-treated lysates subjected to microtubule polymerization and co-sedimentation, probed for EIF4A3 and α-TUBULIN. Each lane was loaded with the entire pellet (P), 1/3 of supernatant containing sample and sucrose cushion (S), and input was 1/10 of total lysate from reaction. Right, agarose gel with rRNA from either mock or RNase A-treated samples. n=5 experiments. **(F)** Cartoon of GST pull down assays for microtubule binding. **(G)** Representative immunoblots of experiments as in **F,** where microtubules were precipitated with GST alone, GST- MAGOH, or GST-EIF4A3 (and probed with anti-GST). Right, GST pull down assay using the RNA and EJC-binding mutant EIF4A3^T163D^. Blot against TUBB3 is shown above. **(H)** Quantification of TUBB3 levels relative to total fusion protein for independent experiments. *p=0.0103 (GST vs. GST-EIF4A3^T163D^), *p=0.0150 (GST-MAGOH vs. GST-EIF4A3^T163D^), **p=0.0.0014 (GST vs. GST- EIF4A3), **p=0.0024 (GST-MAGOH vs. GST-EIF4A3), ns=not significant, one-way ANOVA (Tukey’s multiple comparison test), n=5 (GST), n=6 (GST-EIF4A3) and n=4 (GST-MAGOH and GST- EIF4A3^T163D^) experiments. **(I)** Representative competition experiment in which N2a cells were transfected with FLAG-EIF4A3 with increasing amounts of equimolar V5-MAGOH and V5- RBM8A, and co-IPs of FLAG with endogenous TUBB3. TUBB3/FLAG densitometry ratios: FLAG- EIF4A3 alone normalized to 1 (lane 2), 0.89 (lane 3), 0.42 (lane 4). Arrows, V5-RBM8A (top) and V5-MAGOH (bottom). **(J)** Quantification of competition experiments. *p=0.0261, unpaired t-test (two tailed), n=4 experiments. All graphs, mean + S.D.

To investigate whether EIF4A3 associates with microtubule polymers, we performed microtubule co-sedimentation assays using human cell lysates. Microtubules in lysates were polymerized in the presence of the microtubule stabilizing drug Taxol and then ultracentrifuged onto a glycerol cushion to sediment the polymerized microtubule pellet (**Figure 6D**). Along with α-tubulin, EIF4A3 was enriched in the pellet, indicating it associates with microtubule polymers (**Figure 6E**). Because EIF4A3 is an RNA-binding protein, we tested if RNA cargoes were necessary for EIF4A3 to co-sediment with microtubules by treating cell lysates with RNaseA to degrade single stranded RNA. Co-sedimentation assays using RNA-depleted or untreated lysates showed similar enrichment of EIF4A3 in the microtubule pellet (**Figure 6E**). This indicates that RNA cargoes are not required for EIF4A3 to associate with microtubules.

While these studies demonstrate binding between EIF4A3 and tubulin, they raise the question of whether EIF4A3 and microtubules directly interact. Microtubule co-sedimentation assays with purified GST-EIF4A3 confirmed a direct interaction, which did not occur with GST alone (**Figures S7A**, **S7B** and **S7C**). We further tested this direct interaction by using a GST-affinity pull-down assay with pure microtubules and purified GST, GST-MAGOH, and GST- EIF4A3 (**Figure 6F**). Neither GST alone nor GST-MAGOH associated with microtubules, consistent with co-immunoprecipitation data. In contrast, GST-EIF4A3 did bind microtubules, indicating that EIF4A3 association with microtubules is both specific and direct (**Figure 6G** and **6H**).

We sought to further understand how EIF4A3 associates with microtubules. We first tested if it binds via known EJC- or RNA-binding domains, by using pure proteins of point mutants which impair EJC and RNA binding (GST-EIF4A3^T163D^, EIF4A3^D401E,K402R^ and EIF4A3^E188R^) (Gehring et al., 2009; Ryu et al., 2019). Notably, all 3 mutant proteins bound to microtubules (**Figures 6G**, and **S7D**). This indicates that EIF4A3 can bind microtubules without binding RNA or the EJC.

Given this result, we tested if EJC and microtubule binding to EIF4A3 are independent of each other using competition assays. Increasing amounts of the MAGOH-RBM8A heterodimer reduced tubulin binding to EIF4A3 (**Figure 6I** and **6J**). This experiment demonstrates that association of EIF4A3 with the EJC and microtubules is mutually exclusive. It further suggests that EIF4A3 binding to microtubules may be explained by higher relative levels in the cytoplasm, as we observed (**Figure 5A** and **5B**). Taken altogether with co-immunoprecipitation and co-sedimentation assays, our data demonstrate that EIF4A3 is a bona fide microtubule associated protein (MAP).

With these results, we next turned our efforts to molecular modeling to predict how EIF4A3 specifically associates with microtubules. EIF4A3, like other DEAD-box helicases, can adopt strikingly different conformations depending upon with what it is interacting (Donsbach and Klostermeier, 2021). To determine the most appropriate EIF4A3 conformation, we assessed previously crystalized EIF4A3 structures (**Figure S7E**). These include closed and open conformations of EIF4A3 where it is bound or unbound to MAGOH-RBM8A, respectively (Andersen et al., 2006; Bono et al., 2006; Buchwald et al., 2013). We ultimately selected an open conformation, as co-crystalized with its known interactor CWC22 (PDB 4C9B)(Buchwald et al., 2013), given our findings that MAGOH-RBM8A outcompetes TUBB3 to bind EIF4A3, and that axonal growth is rescued by the EJC- and RNA- mutant EIF4A3^T163D^. For both method validation and as a point of comparison, we also assessed docking and simulation of EIF4A3 and a different conformation of CWC22 to reconstitute this interaction *in silico*, as described in methods.

We sought to define the structure of the tubulin-EIF4A3 complex using protein-protein docking, and molecular dynamics (MD) simulations, and the binding strength using molecular mechanics-generalized Born/surface area (MM-GB/SA) calculations. Application of the docking and filtering procedure to investigate the binding of TUBB3-TUBA1B to EIF4A3 identified two poses (**Figure S7F**). In MD and MM-GB/SA calculations, the binding strength was comparable between TUBB3-TUBA1B-EIF4A3 and CWC22-EIF4A3 (**Figure S7G** and **S7H**). This indicates that EIF4A3 binds tubulin heterodimers with similar binding energies to its known interactors (**Figure 6**). TUBA1B and TUBB3 primarily engaged the C-terminal and N-terminal domain of EIF4A3, respectively (**Figures 7A-C**). The major residues contributing to binding were distributed throughout each of the three proteins **(Figures 7B, 7C, S7I,** and **S7J**). Amongst these, three residues with the largest binding energy, P404, M405, and N406, were adjacent to D401 and E402 residues, which are required for MAGOH-RBM8A interactions (Gehring et al., 2009). This is consistent with a model whereby the MAGOH-RBM8A dimer competes for binding with tubulins. The docking surface of EIF4A3 on TUBB3-TUBA1B corresponds to the microtubule lumen, suggesting that EIF4A3 may bind to both the tubulin heterodimer as well as inner surface of the lattice (**Figure S7K**). Altogether, our molecular modeling supports our discovery that EIF4A3 directly binds to tubulin heterodimers and microtubules.

**Figure 7.**
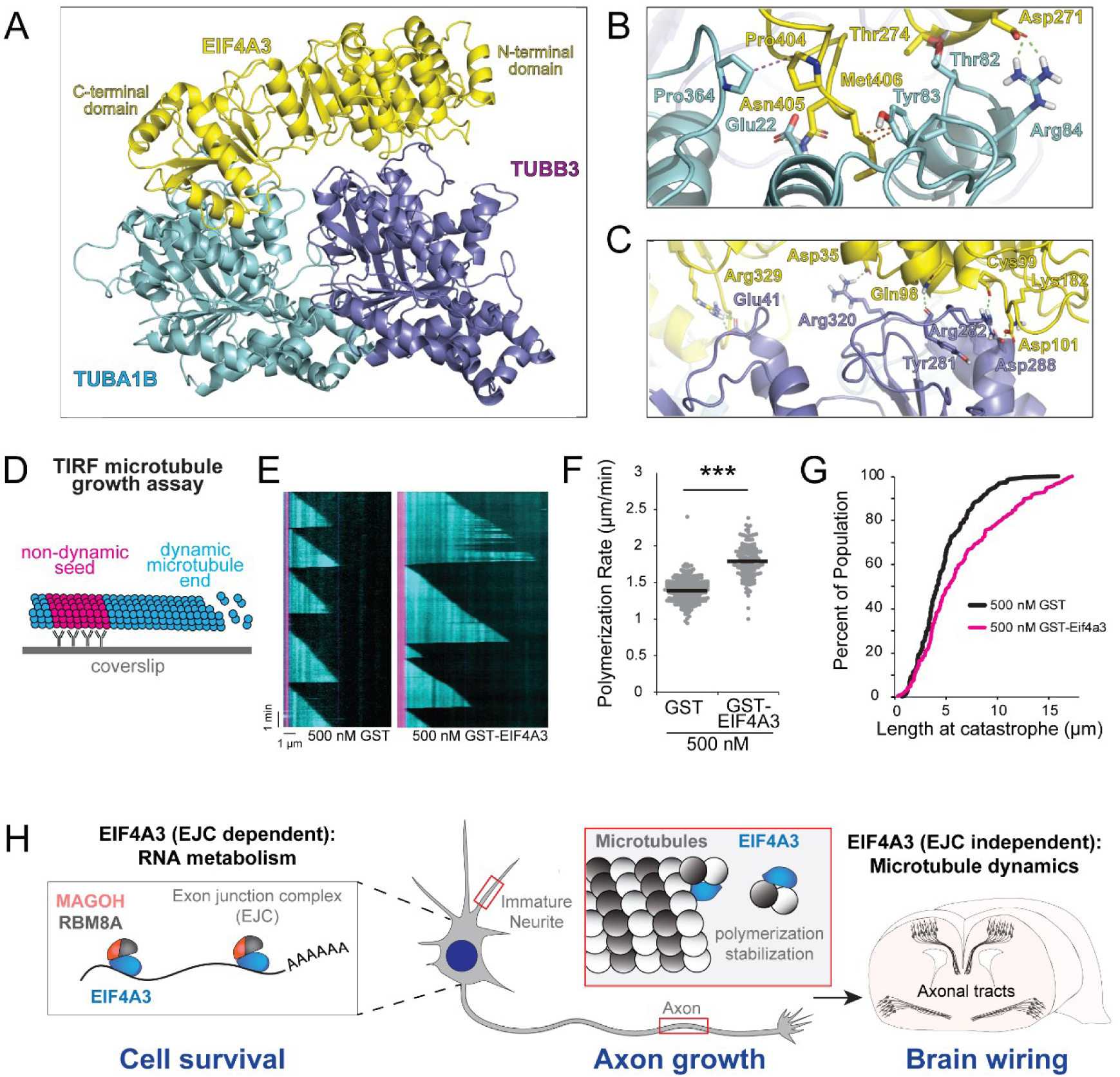
EIF4A3 controls microtubule polymerization and stability. **(A)** Representative structure of TUBB3-TUBA1B docked to EIF4A3 obtained from molecular dynamic modeling. **(B)** Intermolecular interactions taking place between TUBA1B and EIF4A3 in the representative structure. Structure rotated relative to panel A such that TUBB3 is at the back of the view. **(C)** Intermolecular interactions taking place between TUBB3 and EIF4A3 in the representative structure. Structure rotated relative to panel A such that TUBA1B is at the back of the view. In panels B and C, hydrogen bonds (including those as part of salt bridges) are shown with green dashes, CH-π interactions with orange dashes, and CH-CH interactions with pink dashes. **(D)** Cartoon of TIRF (Total Internal Reflection Fluorescence) microtubule growth assay. **(E)** Example kymographs of 15 µM porcine brain tubulin labeled with 10% HyLyte tubulin (cyan) grown from GMPCPP-stabilized seeds (magenta) in the presence of either 500nM GST or 500nM GST-EIF4A3. **(F)** Polymerization rates for microtubules in the presence of 500 nM GST (n=240) or 500 nM GST-EIF4A3 (n=156). Bars, median values. ***p<0.0001, unpaired t-test (two tailed), n=240 microtubules for GST condition (from 4 reactions) and n=156 microtubules for GST-EIF4A3 condition (from 5 reactions). **(G)** Microtubule length at catastrophe plotted as cumulative distributions. **(H)** Graphical summary depicting the dual roles of EIF4A3 (blue): 1) as a canonical RNA-binding protein in the EJC (left) promoting cell survival, and 2) as a microtubule-associated protein promoting microtubule polymerization and stabilization required for axon growth and brain wiring. All graphs, S.D.

### EIF4A3 directly promotes microtubule polymerization and stability

Finally, given that EIF4A3 directly binds microtubules and is required for microtubule stability in neurons, we asked whether EIF4A3 protein alone is sufficient to directly influence microtubule dynamics. To test this, we used Total Internal Reflection Fluorescence (TIRF) microscopy assays (**Figure 7D**) with pure microtubule polymers and either purified GST (control) or GST-EIF4A3. Compared to control, GST-EIF4A3 significantly increased the microtubule polymerization rate by 1.3-fold (**Figures 7E** and **7F**). Further, quantification of microtubule length showed that the presence of EIF4A3 resulted in longer microtubules over time, suggesting that it is protecting microtubules against catastrophe (**Figures 7E** and **7G**). Taken together with the live imaging of cKO neurons, these TIRF findings demonstrate that EIF4A3 is a microtubule binding protein that directly promotes microtubule assembly and stability. Altogether, our findings establish non-canonical RNA-independent functions of EIF4A3 in microtubule regulation which underlies axon growth and brain wiring (**Figure 7H)**.

## Discussion

Our proteome contains ∼1500 RNA-binding proteins, each of which mediate cell-specific and/or housekeeping functions (Baltz et al., 2012; Castello et al., 2012; Gerstberger et al., 2014; Hentze et al., 2018; Van Nostrand et al., 2020). In this study, we challenge the notion that essential RNA-binding proteins exclusively control RNA metabolism. We discover that EIF4A3, the nucleating RNA helicase of the EJC, has been repurposed to directly control microtubule dynamics, independent of its RNA regulatory functions. This non-canonical regulation of the cytoskeleton empowers EIF4A3 to direct axonal growth and brain wiring. Mutations in RNA-binding proteins are abundant in neurodevelopmental and neurodegenerative diseases (Nussbacher et al., 2015; Schieweck et al., 2021), thus, the ability of RNA-binding proteins to also direct microtubule dynamics may have broad implications for disease.

### Dual RNA and microtubule regulators are essential for brain development

EIF4A3 exemplifies an emerging class of dual RNA and microtubule regulators controlling dynamic events of the nervous system. Just a handful of such proteins have been reported, yet this evidence is fragmented (Chudinova and Nadezhdina, 2018). One notable example, Adenomatous polyposis coli (APC), functions as both a microtubule and RNA-binding protein in migrating fibroblasts (Mili et al., 2008; Yasuda et al., 2013). Further, APC is thought to control neuronal migration by directing localization of its mRNA targets at the plus end of microtubules (Preitner et al., 2014). In contrast, our data show that EIF4A3 directly controls microtubules exclusive of its RNA interactions. However, it remains possible that EIF4A3-mediated microtubule regulation ultimately influences RNA metabolism.

RNA-binding proteins may be especially poised to control microtubules given their high cellular abundance, particularly in the cytoplasm. Dual functions for RNA-binding proteins may be particularly pertinent in the nervous system, where rapid control of dynamic activity is needed. For example, proteins like EIF4A3, whose RNA and microtubule functions are in competition, might act as cell sensors to couple gene expression and cytoskeleton growth.

### EJC-independent requirements for EIF4A3 in axonal growth and development

We demonstrate that EIF4A3, but not its canonical EJC partners, is critical for axon growth and *in vivo* formation of long-range intracortical axonal tracts. This is relevant for neurodevelopmental disorders in which aberrant long-range underconnectivity is implicated (Kern et al., 2015). We generated, for the first time, RCPS patient-derived cortical organoids, showing that *EIF4A3* mutations also cause neuronal maturation defects. These data suggest that defective neuronal maturation may contribute to language and learning impairments in RCPS patients, as well as intellectual disability associated with *EIF4A3* mutation.

Beyond axon formation, EIF4A3’s noncanonical functions may additionally underlie neuronal processes reliant upon dynamic microtubule control, such as dendrite and synapse function (Aiken and Holzbaur, 2021; Barker-Haliski et al., 2012; Giorgi et al., 2007). Microtubule properties differ substantially over the course of neuronal development (Lasser et al., 2018; Penazzi et al., 2016; Song and Brady, 2015). In particular, dendrite and axonal microtubules diverge in their directionality of growth, interactions with microtubule-associated proteins, and tubulin post-translational modifications. Thus, RNA- and microtubule-dependent requirements of EIF4A3 might also be crucial during later stages of neuronal maturation.

### EIF4A3 is a non-canonical microtubule-associated protein

Altogether, our data argue that EIF4A3 promotes neuronal survival in an EJC-dependent fashion and independently directs dynamic events of axonal growth via microtubules. We find that cytoplasmic pools of EIF4A3, which are not bound to the EJC and RNA, promote microtubule stability and polymerization. This mutually exclusive binding to the EJC or microtubules is evidenced by both competition and sub-cellular localization experiments. It is also consistent with proteomic data of the developing brain which report more EIF4A3 relative to other core EJC components (Mao et al., 2016a). EIF4A3 requires an intact EJC to stably bind RNA (Andersen et al., 2006; Bono et al., 2006) and thus microtubule-bound EIF4A3 may be unlikely to recruit RNA. Going forward it will be illuminating to elucidate how the switch of EIF4A3 binding from EJC to microtubules is achieved. Indeed, the EJC is a dynamic multiprotein complex (Le Hir et al., 2016; Mabin et al., 2018) and distinct binding partners may enhance an EJC- or microtubule-bound conformation. In the presence of MAGOH-RBM8A and RNA, EIF4A3 is in a closed conformation (Bono et al., 2006), which may prohibit microtubule binding. This RNA-microtubule switch might also be achieved by differential subcellular localization and post-translational modifications.

Our data indicate that EIF4A3 is a microtubule-stabilizing protein. The *in vitro* reconstitution experiments demonstrate that EIF4A3 promotes microtubule polymerization and suppresses catastrophe. EIF4A3’s impact on microtubules resembles functions of microtubule-stabilizing proteins such as Tau and p150^Glued^ (Drechsel et al., 1992; Lazarus et al., 2013). Whereas Tau binds to microtubules and alters lattice architecture by increasing protofilament number (Prezel et al., 2018), p150^Glued^ binds tubulin dimers and microtubules, and is thought to regulate microtubule dynamics at plus ends (Ligon et al., 2003). Our biochemical studies suggest that EIF4A3 can bind both microtubule polymers and tubulin heterodimers. Molecular modeling is consistent with three possible binding configurations whereby EIF4A3 may associate with tubulin heterodimers, the plus-ends, and/or along the inside of the microtubule lattice. We thus favor a model in which EIF4A3 associates with α- and β-tubulin heterodimers to promote microtubule growth and is retained at the growing ends of microtubules (**Figure 7H**). The binding to the inner part of the lattice is intriguing and reminiscent of microtubule inner proteins (MIPs) that have been identified in cilia but are poorly understood (Ma et al., 2019). In future studies, it will be important to understand the biophysical properties by which EIF4A3 controls microtubule polymerization and stability, as well as the relative contribution of predicted EIF4A3- tubulin interacting residues.

### The exon junction complex and unique contributions to disease

Our study shows definitively that the central RNA-regulator EIF4A3, can function non-canonically in an EJC- and RNA-independent manner. EJC components, and in particular EIF4A3, are implicated in diverse cellular processes, including cell division, RNA localization, and cell fate (Akhtar et al., 2019; Budiman et al., 2009; Ishigaki et al., 2014; Kanellis et al., 2021; Kwon et al., 2021; Pilaz et al., 2016; Wang et al., 2021). We propose that beyond the nervous system, EIF4A3 may contribute to these cellular processes via microtubule regulation. Although previous studies have suggested that EJC components can function outside the complex (Alexandrov et al., 2011), in all cases these are RNA-dependent functions, and EJC independency has never been shown. Thus, our findings highlight the complexity of the EJC and the need for further functional studies within and outside the complex.

The three core EJC components are broadly expressed yet are implicated in divergent disorders and cellular processes. Although *RBM8A* and *EIF4A3* copy number variations are both associated with intellectual disability, their loss of function mutations cause vastly different developmental disorders- TAR syndrome and RCPS (Albers et al., 2013; Bertola et al., 2017; Favaro et al., 2014). Our study brings novel insights into this apparent paradox, and we speculate that some clinical differences may be explained by EIF4A3 EJC-independent functions. More broadly, EIF4A3’s non-canonical microtubule function may also be relevant for cancer, in which EIF4A3 is also implicated (Ye et al., 2021). In sum, our study generates a new framework for how an indispensable RNA regulatory complex can control physiology within and outside the nervous system.

## Acknowledgements

We thank Silver lab members for helpful discussions as well as Cagla Eroglu, Josh Huang, Dong Yan, and Terry Lechler for careful reading of the manuscript. We thank Emily Miller for co-sedimentation assays; Ashley Lennox for initial analysis of *Eif4a3* cHets; Jeremy Rouanet and Charlie Sheehan for technical assistance. We are grateful to the following for reagents and equipment: Yingwei Mao (*Rbm8a* floxed mice), Klaus Nave (*Nex*-Cre mice); Santos Franco and Jonathan Cooper (*Dcx*-Cre plasmids), Maria Rita Passos-Bueno (*RCPS* iPSCs), Vann Bennett (GST plasmid), Tom Petes and Vann Bennett (ultracentrifuges), the Duke FACs facility, and Duke Light Microscopy Core Facility. The following grants supported this research: to DLS (NIH: R01NS083897, R01NS120667, R01NS110388), to Duke Light Microscopy Core Facility (S10OD020010), to FA (Ruth K Broad fellowship, International Brain Organization (IBRO) fellowship), to MA (Raine Priming Grant, Raine Medical Research Foundation). To JM (NIH: R35GM136253).

## Author contributions

FA and DLS conceived of the project and wrote the paper. DLS oversaw research. FA performed analyses of EJC mutant brains, with the help of FM and BL. FA performed *in utero* electroporations, microtubule live imaging, co-immunoprecipitations and FACS. FA and BL performed SIM and STED imaging. BL performed GST pulldowns and purifications. LL performed CUBIC experiments with the help of FA. FA and BL performed primary neuronal cultures, with the help of CN for data analysis. CM and FA performed cortical organoid experiments with the help of FM. LW and JM performed TIRF experiments. MA performed molecular modelling experiments.

## Declaration of interests

The authors declare no competing interests.

## STAR Methods

### Mouse husbandry and genetics

All animal procedures were approved by the Duke Institutional Animal Care and Use Committee (IACUC) and performed in agreement with the ethical guidelines of the Division of Laboratory Animal Resources (DLAR) from Duke University. We used the following previously described mouse lines: *Eif4a3*^loxP^ (Mao et al., 2016b), *Rbm8a*^loxP^ (McSweeney et al., 2020), *Magoh*^loxP^ (McMahon et al., 2014) and *Neurod6^tm1(cre)Kan^* (*Nex*-Cre) (Goebbels et al., 2006). The following mouse strains were obtained from Jackson Laboratories: C57BL/6J (*wild type*), B6.Cg-Gt(ROSA)26Sor^tm14(CAG-tdTomato)Hze^/J (Ai14), B6.129S2-Trp53^tm1Tyj^/J (p53^LoxP^), and C57BL/6J- Tg(Dcx-DsRED)14Qlu/J (*Dcx*::DsRed). For embryo staging, plug dates were defined as embryonic day (E) 0.5 on the morning the plug was identified.

### Cell lines and mouse primary neuronal cultures

HeLa and N2a immortalized cell lines were maintained at 37°C in 5% CO2 environment, with DMEM high glucose media (Gibco™) supplemented with 10% fetal bovine serum (HyClone) and 1% Pen-Strep (Gibco™). For primary cultures, mouse embryonic cortices were dissected at the specified stage and digested at 37°C for 10 min with 5 U/ml papain (Sigma) and 120 U/ml DNase I (Worthington) in PBS supplemented with 0.25 mg/ml DL-cysteine HCl (Sigma), 0.25 mg/ml BSA (Sigma) and 6.25 mg/ml D-glucose (Sigma). Then, cortices were rinsed twice with Neurobasal media (Gibco™) and mechanically triturated with the pipette in neuronal media (see below) supplemented with 120 U/ml DNase I. The dissociated neurons were counted, and 1.75 x 10^5^ cells were seeded in wells of 24-well plates with 12 mm coverslips (Carolina® 0.09-0.12 mm thickness) pre-treated with acetone and coated with 0.1 mg/ml poly-D-lysine (Sigma; 30,000- 70,000 mol wt). Neuronal media recipe: Neurobasal (Gibco™), 1% Glutamax (Gibco™), 2% B27 (Gibco™) and 0.12% Pen-Strept (Gibco™). Axonal length and neurite outgrowth were quantified using the NeuronJ plugin from Fiji (ImageJ). Axons were identified morphologically as the longest projection of polarized neurons (stage 3). For rescue experiments, CAG-GFP was co-electroporated to visualize and assess neuronal morphology.

### *In utero* electroporation (IUE)

*In utero* electroporation was performed as previously reported (Saito and Nakatsuji, 2001). Briefly, E14.5 or E15.5 pregnant females were anesthetized with isoflurane. Uterine horns were exposed by making an incision in the abdomen. Each embryo was injected with 1-1.5 ul of plasmid solution (containing 0.01% fast green and 1-3 μg/μl of plasmids) and electroporated using the following parameters: five 50 ms-pulses at 45V (E14.5) or 50v (E15.5) with 950 ms pulse-interval, using platinum-plated BTX Tweezertrodes. Then, uterine horns were repositioned into the abdominal cavity and the muscle and skin incisions were sutured. IACUC procedures were followed to ensure appropriate care and anesthesia of the animals.

### Electroporation of primary neuronal cultures and cell lines

For transfection of primary neuronal cultures, the standard protocol for the Lonza® P3 primary cell 4D Nucleofector™ X Kit S was followed using pulse code CA-138, right after dissection and single cell suspension. For the electroporation, 1 x 10^6^ cells and 2 ug plasmid DNA (1:4 pCAG-GFP:shRNA ratio) were used for each well in the 16-well cuvette. For transfection of N2a cells, Lipofectamine® 2000 transfection reagent (Invitrogen) was used according to manufacturer’s protocol. Briefly, cells were split the previous day to get ∼60-80% confluency the day of transfection, and were transfected with 7.5 μl of lipofectamine and 3 μg of DNA per well of a 6- well plate (for a 60 mm plate, 10 ul of lipofectamine and 5 μg of DNA were used). The following shRNAs were used from Open Biosystems/Horizon: pGIPZ non-silencing lentiviral shRNA control, pGIPZ shRNA *Eif4a3* (AGGTGTCCCTCATCATTAA), pGIPZ shRNA *Magoh* (TTGTAATTGCTGTTGTTGG); and Sigma: pLKO.1 shRNA *Rbm8a* (CCGGGAAGGACTAAATGGTCAAGATCTCGAGATCTTGACCATTTAGTCCTTCTTTTTG).

### Immunofluorescence staining and image acquisition

Cell cultures were fixed for 15 min at room temperature with 37°C pre-warmed 4% paraformaldehyde (PFA) (Sigma) in PBS. Then rinsed with PBS and permeabilized with 0.5% Triton X-100 (Sigma) in PBS for 10 min. Primary and secondary antibody incubations and blocking were done in 10% normal goat serum in PBS. Washes were done with 0.1% Tween 20 detergent (Bio-Rad) in PBS. For tissue sections, brains were fixed overnight in 4% PFA-PBS at 4°C, rinsed in PBS and followed by submersion in 30% sucrose-PBS until sinking (24-48 hrs). Brains were frozen in NEG-50 medium (Richard-Allan Scientific) and cryostat sections (20 μm) were prepared and stored at -80°C until use. Sections were permeabilized with 0.3% Triton X-100 for 10 min and blocked with 5% normal goat serum in PBS for 1 hr. at room temperature. Sections were incubated with primary antibodies overnight at 4°C. Then they were washed 3 times 10 min with PBS and incubated in species appropriate secondary antibodies and 1 ug/ml Hoechst 33342 (Invitrogen) or DAPI (Invitrogen) for 30 min at room temperature. Finally, they were washed as described before and mounted. Mowiol® (Sigma), Fluoromount-G™ (Invitrogen) or Vectashield® (Vector Laboratories) were used as mounting media. Images were acquired with a Zeiss Axio Observer Z.1 microscope coupled with an apotome2, unless otherwise specified. Image measurements and quantifications were blindly performed using Fiji (ImageJ). A minimum of 3-6 sections from anatomically comparable regions per embryo were analyzed on each case, coming from at least 3 biological replicates from control and mutant littermate embryos of at least 2 independent litters. For L1+ axonal tract thickness, sections from similar anatomical regions in the motor/somatosensory cortex from littermate mice were analyzed, and the same threshold was applied for all images from all experimental conditions. Apoptotic cells were identified as CC3+ cytoplasmic staining with pyknotic nuclei. For quantification of the signal of EJC components in nucleus and axon, the fluorescence intensity was quantified by the following formula: Corrected total cell fluorescence (CTCF) = Integrated density – (Area of selected cell region X Mean grey fluorescence of background readings). Briefly, the nucleus and the first 10 μm of the axon were selected using the polygon selection tool of ImageJ and integrated density and area measurements were obtained. For the background 3 random measurements of a 7 x 7 μm square was obtained from each image.

### Super-resolution microscopy

For STED microscopy, neuronal cultures were prepared from E15.5 mice onto #1.5 cover slips and fixed at DIV2 as described above. Prolong glass mounting media was used. Cell nuclei were stained using Quant-IT™ PicoGreen™ dsDNA reagent (ThermoFisher Scientific; 1:1000). Images were acquired in a Leica DMi8 STED and confocal microscope from Duke Light Microscopy Core Facility.

### Antibodies

The following primary antibodies were used: L1CAM (Millipore MAB 5272; 1:500 IF), CC3 (Cell Signaling 9661; 1:400 IF), β3-TUBULIN/TUBB3/TUJ1 (BioLegend 801202; 1:2000 WB or 1:1500 IF), β3- TUBULIN/TUBB3/TUJ1 (Cell Signaling D71G9 [5568]; 1:500 WB or 1:1000 IF), GFP (Abcam ab13970; 1:1000 IF), EIF4A3 (Bethyl A302-980A; 1:500 IF), EIF4A3 (Abcam ab180573; 1:500 IF or 1:1000 WB), EIF4A3 (Santa Cruz sc-67369; 1:50 WB), EIF4A3 (Abcam ab32485; 1:300 IF or 1:900 WB), MAGOH (Proteintech 12347-1-AP; 1:100 IF), MAGOH (Santa Cruz sc-56724, 1:50 WB), RBM8A (Santa Cruz sc-32312; 1:100 IF), RBM8A (Proteintech 14958-1-AP; 1:50 WB), α-TUBULIN (Sigma T6199; 1:1000 WB), FLAG (Sigma F1804; 1:400 IF or 1:500 WB), V5 (Sigma V8137; 1:1500 WB), V5 (Cell Signaling 13202 [D3H8Q]; 1:1000 WB), GST (Santa Cruz sc-138; 1:1000 WB), SOX2 (ThermoFisher Scientific 14-9811-80; 1:500 IF), PAX6 (Biolegend 901301; 1:1000 IF), p53 (Leica CM5; 1:500 IF), ANKYRIN-G (NeuroMab 75-146, clone N106/36; 1:100 IF), SMI-312 (Biocompare 837904; 1:500 IF).

The following secondary antibodies were used: Alexa Fluor 488 (Invitrogen; 1:800), Alexa Fluor 594 (Invitrogen; 1:800), Alexa Fluor 647 (Invitrogen; 1:800), for STED: ATTO 647N (Active Motif; 1:500), goat anti-mouse HRP conjugated (ThermoFisher Scientific; 1:5000), goat anti-rabbit HRP conjugated (ThermoFisher Scientific; 1:5000), goat anti-mouse light chain specific HRP conjugated (Jackson ImmunoResearch Laboratories; 1:5000), mouse anti-rabbit light chain specific HRP conjugated (Jackson ImmunoResearch Laboratories; 1:5000).

### Tissue clearing (CUBIC)

Whole-mount embryonic heads were cleared using an adapted version of CUBIC protocol(Susaki et al., 2015). Briefly, samples were fixed in 4% PFA-PBS overnight at 4°C and then incubated in CUBIC ½ R-1 (1:1 dilution of CUBIC R-1 solution in dH2O) overnight at 37°C for up to 3 days depending on the sample size. Samples were then incubated in CUBIC R-1 solution (25 wt% urea, 25 wt% Quadrol, 15 wt% Triton X100, dH2O) for 1-3 days at 37°C with rocking until completely clear. Afterward, samples were incubated in CUBIC ½ R-2 (1:1 dilution of CUBIC R-2 in 1x PBS) overnight at 37°C until sinking and then finally incubated in CUBIC R-2 solution (25 wt% urea, 50 wt% sucrose, 10 wt% triethanolamine, dH2O) solution at 37°C until clear and ready for imaging. Between incubation in CUBIC R-1 and CUBIC ½ R-2, samples were washed in PBS/0.01% NaN3 at room temperature with rocking to remove excess CUBIC R-1 to improve clearing efficiency. Once cleared, samples were mounted and imaged using a 5x objective of a Zeiss Lightsheet Z.1 microscope from Duke Light Microscopy Core Facility. 3D reconstructions were generated and quantified with Imaris software. As specified, IUE was combined with the adapted CUBIC tissue clearing protocol to visualize sparsely labelled neurons in embryonic mouse brains at single-cell resolution. In this case, embryonic mouse brains were specifically isolated to improve clearing and obtain single-cell resolution. Cleared samples were imaged as described above using both 5x and 20x objectives. 3D reconstructions were then quantified and compared by measuring the length of axonal tracts per brain, which was defined as the average length of the five longest axons found in the IUE region. The volume of td-tomato fluorescence specifically in axonal tracts found within the IUE region was also measured and compared between samples. Fluorescence volume in axonal tracts was evenly measured across samples with different sized IUE regions by measuring a region of set size and volume across all images. All quantifications were conducted blind, and 3 brains were quantified for each genotype across 2 different mouse litters. Statistical significance was determined using one-tailed Student’s t-tests.

One-tailed Student’s t-tests were used due to our expectation that axonal length and volume would be significantly reduced based on our prior data.

### Single molecule fluorescence in situ hybridization (smFISH)

*Mus Musculus Eif4a3* smFISH probes were designed and prepared as previously described (Tsanov et al., 2016). For *in situ* hybridization, tissue sections were permeabilized in 0.5% Triton X-100 in PBS for 30 min at room temperature and rinsed two times with wash buffer (10% formamide, 2x SCC buffer). Then, smFISH probes were diluted 1:200 in hybridization buffer (10% formamide, 2x SCC buffer, 10% dextran sulfate) and incubated overnight at 37°C. Finally, samples were washed two times at 37°C (Hoechst was included in second wash). Vectashield was used as mounting media. To quench RNase activity, all buffers were prepared with diethyl pyrocarbonate (DEPC) water.

### Western blot and co-immunoprecipitations

Cell cultures were lysed at 4°C in TNE buffer (50 mM Tris-HCl pH=7.4, 137 mM NaCl, 0.1 mM EDTA) containing 0.5% Triton X-100, plus protease inhibitors (Roche). Protein lysates were incubated on ice for 10 min and clarified by centrifugation at 20,000 rcf for 5 min. For *in vivo* tissue co-immunoprecipitations, mouse cortices from 6-8 E16.5 C57BL/6J embryos were homogenized in lysis buffer using a PYREX® tissue grinder. For co-immunoprecipitations 1% octyl-beta-glucoside was added to the lysis buffer, and samples were incubated for 40 min. on ice and clarified by manual spin for 15 sec. at 4°C. 0.5-1 ml of each protein lysate was incubated with 5 ug/ml of primary antibody overnight at 4°C with shaking. The following day protein G magnetic beads (Bio-Rad or Invitrogen) were prepared according to the manufacturer’s protocol and samples (protein lysate + primary antibody) were added to the beads and incubated for 2 hrs. at 4°C or 1 hr. at room temperature with shaking. Then beads were magnetized and after 5 washes with TNE buffer plus 0.5% NP-40 detergent and protein inhibitors, beads were resuspended in elution buffer (TNE buffer plus 1x laemmli buffer and 50 mM DTT) and boiled 5-10 min. at 95°C. For Western blot, 20-50 ug of protein lysates, plus 1x laemmli buffer and 50 mM DTT, were boiled for 5 min. at 95°C. SDS-PAGE was run using 4-20% Mini-PROTEAN® TGX™ precast gels (Bio-Rad) and run at 100-150V for 1-2 hrs. Blotting was done using the Trans-Blot Turbo Transfer System (Bio-Rad). Blots were all blocked in 5% milk in TBST (TBS buffer plus 0.1% tween detergent). Primary antibodies were incubated overnight at 4°C shaking. The following day, they were rinsed 5 times 15 min. in TBST buffer, incubated with secondary antibody in TBST buffer for 1 hr. at room temperature and rinsed 5 times 15 min. in TBST buffer. Western imaging was done using ECL Western blotting substrate (Pierce™) supplemented occasionally with SuperSignal™ West Femto Maximum Sensitivity Substrate (Thermo Scientific). Blots were exposed in a Bio-Rad Gel Doc XR system. Quantification of Western Blot was performed using ImageJ software with values normalized to α-TUBULIN loading controls as well as background. In co-immunoprecipitations from cortical tissue either FLAG antibody or RFP antibody were used as control (same species as TUBB3 and EIF4A3 antibody, respectively).

### RT-qPCR and primers

RNA was extracted in RLT buffer plus 0.01% β-mercaptoethanol following RNeasy kit (QIAGEN) and cDNA was prepared using the iScript kit (Bio-Rad) following manufacturer’s protocol. qPCR was performed using Sybr Green iTaq (Bio-Rad) in, at least, 3 independent biological samples (3 technical replicates per sample) in a QuantStudio 3™ machine (Applied Biosystem™). Values were normalized to *β-Actin* as loading control. The following primers were used: *Mus Musculus β-Actin* (Forward 5’- AGATCAAGATCATTGCTCCT -3’ and Reverse 5’- CCTGCTTGCTGATCCACATC -3’), *Mus Musculus Eif4a3* (Forward 5’- -3’ and Reverse 5’- -3’), *Mus Musculus Rbm8a* (Forward 5’- GCTCTGTTGAAGGTTGGATTCT -3’ and Reverse 5’- GCCCCATCAAATCTTGACCAT -3’), *Mus Musculus Magoh* (Forward 5’- ACTTTTACCTGCGTTACTACGTG -3’ and Reverse 5’- GTTGTTGGCGTATCGCAATTT -3’), *Mus Musculus MagohB* (Forward 5’- AGTTTTTGGAGTTCGAGTTTCGG -3’ and Reverse 5’- AATGTGTTCATCCCCAATTACGA -3’)(Spandidos et al., 2010), *Homo sapiens TBP* (Forward 5’- GTGACCCAGCATCACTGTTTC -3’ and Reverse 5’- GCAAACCAGAAACCCTTGCG -3’), *Homo sapiens OCT4* (Forward 5’- GTGGTCAGCCAACTCGTCA -3’ and Reverse 5’- CCAAAAACCCTGGCACAAACT -3’), *Homo sapiens NANOG* (Forward 5’- TGGACACTGGCTGAATCCTTC -3’ and Reverse 5’- CGTTGATTAGGCTCCAACCAT -3’), *Homo sapiens SOX17* (Forward 5’- GTGGACCGCACGGAATTTG -3’ and Reverse 5’- GGAGATTCACACCGGAGTCA -3’), *Homo sapiens BRACH* (Forward 5’- TATGAGCCTCGAATCCACATAGT -3’ and Reverse 5’- CCTCGTTCTGATAAGCAGTCAC -3’), *Homo sapiens PAX6* (Forward 5’- TGGGCAGGTATTACGAGACTG -3’ and Reverse 5’- ACTCCCGCTTATACTGGGCTA -3’), *Homo sapiens FOXG1* (Forward 5’- AACCTGTGTTGCGCAAATGC -3’ and Reverse 5’- AAACACGGGCATATGACCAC -3’), *Homo sapiens EIF4A3* (Forward 5’- GGAGATCAGGTCGATACGGC -3’ and Reverse 5’-GATCAGCAACGTTCATCGGC -3’).

### Fluorescence activated cell sorting (FACS)

GFP+ or tdTomato+ cells were collected in 10% fetal bovine serum in ice-cold HBSS buffer (Gibco™) plus 2 U/ml DNase I RNase free (NEB®) and 1.25 ug/ml propidium iodide (Invitrogen) or 5 ug/ml DAPI, respectively. Then, samples were filtered with 30 um cell strainers into sorting tubes, and samples were sorted using 488 nm or 561 nm lasers of a B-C Astrios Sorter from the Duke Cancer Institute flow cytometry facility. Samples were collected directly in RLT buffer plus 1% β-mercaptoethanol (Sigma) and RNA was extracted with the RNeasy kit (QIAGEN), according to manufacturer’s protocol. For mouse brains, embryonic cortices were dissected and dissociated in 0.25% trypsin-EDTA (Gibco™) for 10 min. at 37°C and processed as specified above.

### Live imaging

*Eif4a3*^+/+^ or *Eif4a3*^lox/lox^ littermate mouse embryos were co-electroporated with GFP-Macf43 (also known as GFP-MT+TIP) (Honnappa et al., 2009; Yau et al., 2016) and *Dcx*-mCherry-IRES-Cre (gift of Francos Santos, Jonathan Cooper) at E14.5. The following day, electroporated cortices were dissected and neuronal cultures were prepared as described above. Neurons were plated in 24-well glass bottom plates and DIV2 neuronal cultures were imaged using a Leica DMi8 Andor Dragonfly Spinning Disk Confocal plus microscope from Duke Light Microscopy Core Facility. Cells were maintained in incubation chamber at 37°C and 5% CO2. Imaging conditions included 63x oil objective, 2x camera zoom, 5% laser intensity, 300 ms exposure time and 500 gain. Neurons were randomly selected, checked for red fluorescence in 594 channel (for the presence of Dcx-mCherry-IRES-Cre) and then imaged using the 488 channels for 1 minute (every 500 ms). To trace each particle, the “segmented line” tool was used to manually select the neurites to be analyzed, and the “multiple kymograph” plugin from ImageJ was used to generate kymographs that included the movements of each comet across time. Uninterrupted movements were manually traced with the “straight line” tool and the “bounding rectangle” measurement was extracted to obtain angle and length and calculate the distance, lifetime and speed based on scale. Microtubule catastrophe frequency is expressed as the inverse of the average lifetime or distance of all the comets in each neurite. Thousands of comet movements coming from multiple neurites (n=1-6 per cell), neurons (n=15 per genotype) and embryos from different litters (n=3 per genotype).

### Cloning

To generate the GST-EIF4A3 used in the co-sedimentation assays, *Homo sapiens EIF4A3* cDNA was cloned into a pGEX-6P-1 6xHis pDEST GST vector using Gateway® cloning technology (ThermoFisher Scientific). To generate *Mus Musculus* 3xflag-EIF4A3 and 3xflag-MAGOH, 3xFLAG tag was generated by Genscript and put into pUC57 flanked by EcoRI and XbaI. This plasmid was digested and put into a linearized pCAG vector thus generating pCAG 3xFLAG. Full-length *Eif4a3* was amplified from *Mus Musculus* cDNA using primers that contained XbaI and XhoI sites. The TOPO-TA kit was used to put the insert into a plasmid and then digested and gel extracted. pCAG 3xFLAG vector was linearized using XbaI and XhoI and a standard T4 DNA ligation reaction was done and transformed into competent cells. Point mutants (401-402, 188 and T163D) were made by Gibson Assembly® (NEBuilder® HiFi DNA Assembly) according to manufacturer’s protocol, using EcoRI-HF and SapI cut sites in the pCAG 3xFLAG vector. We generated equivalent amino acid substitutions as previously described for *Homo Sapiens EIF4A3* mutants(Gehring et al., 2009; Ryu et al., 2019): GACGAG (D401-E402) to AAGCGC (K401-R402) for EIF4A3 401-402 mutant; GAG (E188) to CGC (R188) for EIF4A3 188 mutant; and ACG (T) for GAC d, for EIF4A3 T163D mutant. To generate GST-MAGOH, GST-EIF4A3 and GST-EIF4A3 truncated mutants, NEBuilder® HiFi DNA Assembly was followed, using NotI and EcoRV cut sites in the pCAG 3xFLAG vector and then subcloned into a pGEX-6P-1 6xHis pDEST GST vector.

### GST protein purification

To purify GST tagged EIF4A3 constructs, GST tagged MAGOH and GST alone, BL21 competent cells were transformed, and one colony was picked to grow a 5 ml culture in 2x YT media overnight at 37°C shaking at 250 rpm. The next morning, bacteria culture was transferred into 300 mL of 2x YT media and allow to grow for 2 hrs. at 37°C, shaking at 250rpm. At 2 hrs. post-inoculation, 1 µM IPTG and 2% glucose were added to the culture, and cultures were allowed to grow for an additional 4 hr. at 30°C and 160rpm. Bacteria were then collected and brought up in standard buffers with protease inhibitors (Roche) according to the standard protocol for batch purification with the Pierce™ Glutathione Agarose (ThermoFisher Scientific) or GST Bulk Kit (GE Healthcare). Samples were then sonicated 10 sec. on, 60 sec. rest, 10 times at 30% amplitude on ice. Lysates were clarified by spinning at 4°C for 20 min at 3220xg. To purify the protein, clarified lysate was incubated with glutathione (GSH) agarose beads for 1 hr. at room temperature, centrifuged 2 min 700 rcf, washed 3 times in wash buffer (50mM Tris, 150mM NaCl, pH=8) and eluted with 10 mM reduced glutathione (ThermoFisher Scientific) in wash buffer three successive times.

### GST pull-down assays

Protocol for GST purification was followed as described above and then stopped before the elution. Then, after the first series of 3 washes, 20 ug (0.4 nmol) of purified microtubules (Cytoskeleton) were added to the beads and incubated for 30min at room temperature. Samples were washed 3x and samples were eluted with 1X reduced glutathione, three successive times. Binding of microtubules to the GST-tagged protein was determine by presence of the former in the eluate. Eluates were loaded onto 12% Mini-PROTEAN TGX gels and SDS-PAGE and Western analysis were performed as previously described. As this assay was performed at room temperature but in the absence of taxol, it would detect interactions which may include free tubulin and polymers.

### Microtubule co-sedimentation assays

The standard protocol for Microtubule Binding Protein Spin-down Assay Kit was used according to manufacturer’s protocol (Cytoskeleton). Briefly, microtubules were polymerized at 35°C and incubated with purified GST-EIF4A3 for 30 min at RT. Cellular lysates were collected in RIPA buffer (Pierce™) supplemented with a protease inhibitor cocktail (Roche) and protein quantity was ascertained using a standard BCA assay (Pierce™). For RNase treatment of microtubule pulldowns lysates were subjected to 0.15mg/mL RNaseA for 30 min at RT. RNA was isolated from these input samples with TriReagent (Sigma) using their standard protocol. rRNA samples were run on a 1% DEPC agarose gel to assess RNA quality. As specified by manufacturer, 100 μl of glycerol cushion buffer and 50 ul including taxol polymerized microtubules + fusion protein or lysate were used per condition. Ultracentrifugation steps were performed using the Beckman Coulter TLA120.1 rotor and were spun at 100,000g. For SDS-PAGE analysis one-third of the supernatant containing sample and glycerol cushion were run, reducing the yield of detectable protein in the supernatant.

### Brain cortical organoids generation, maintenance and dissociation

Work with iPSCs was approved as a human exempt by the Duke University Institutional Research Board. The iPS cell lines (identity is blind to the authors) used in this study were generated previously characterized and genotyped(Miller et al., 2017). Cells were maintained in Essential 8™ medium (Gibco™) supplemented with 100 ug/ml Normocin™ (Invivogen) on Matrigel (Corning™)-coated plates, and then seeded on vitronectin (Gibco™) for 2 passages before generating the 3D cultures. Cortical organoids were generated and maintained as previously described(Yoon et al., 2019). Dissociation of organoids in 2D cultures were performed as previously described (Miura et al., 2020) with minor modifications. The following lines were dissociated: 8799, 7405, 8417-1, 8417-2. Thus, 5 x 10^4^ cells were seeded in wells of 24-well plates with 12 mm coverslips pre-treated with acetone and coated with poly-D-lysine as described above for mouse primary neuronal cultures. Cultures were maintained in the same medium used for >D43 organoids and fixed for 15 min at room temperature with 37°C pre-warmed 4% paraformaldehyde (PFA) in PBS. AnkG puncta and NeuN+ cells were blindly and automatically quantified by QuPath software. To generate an isogenic control line, we corrected the mutation in a RCPS patient-derived iPSC line (F8417-2cl3) using CRISPR/Cas9 genome editing. The disease-associated 16 repetitive motifs in one allele of the *EIF4A3* 5’UTR in the patient line was replaced with 7 motifs to obtain a heterozygous isogenic control. The control donor motifs used included 4 CACA-20nt, 1 CA-18nt, 1 CACA-20nt, and 1 CA-18nt (see Figure S4B); and the whole genome sequence that includes them is the following: TCGGCAGCGGCACAGCGAGGTCGGCAGCGGCACAGCGAGGTCGGCAGCGGCACAGCGAGGTCGGCAGCGGCACAGCGAGGTCGGCAGCGGCAGCGAGGTCGGCAGCGGCACAGCGAGGTCGGCAGCGGCAGCGAGG. Plasmid pX330 encoding two guide RNAs and Cas9 (2μg) and plasmid pCAG-EX2 with a homology repair donor sequence and Puromycin resistance cassette (4μg) were electroporated into 2x106 cells using the P3 Primary Cell 4D-Nucleofector X Kit L (Lonza, V4XP-3024). Following electroporation, cells were seeded across 6 wells in a Matrigel-coated 6-well plate in Essential 8 medium supplemented with 10μM ROCK inhibitor (StemCell Technologies). Cells were selected with Puromycin (0.25μg/ml) beginning 24h after electroporation for 6 days. Single iPS colonies were isolated, expanded, and screened by PCR using genotyping primers (Forward 5’- ACGCCCAGTTCCCTTTCAC -3’ and Reverse 5’- GAACGTGGGGGTCACATC -3’). The selected edited line was further characterized by Sanger sequencing of EIF4A3 5’UTR, RTqPCR and immunofluorescence of pluripotency markers before generating cortical organoids.

### *In vitro* Microtubule Dynamic Assays

Microtubule seeds were assembled by incubating 21.6 µM porcine brain tubulin containing 23% rhodamine-labeled tubulin (Cytoskeleton, Inc; Denver, CO) in BRB80 buffer (80 mM PIPES brought to pH6.9 with KOH, 1 mM ethylene glycol tetraacetic acid (EGTA), 1 mM MgCl2 with 1 mM GMPCPP (Jena Biosciences; Jena, Germany) at 37°C for 30 minutes. Sample was then centrifuged at 126,000 x g for 10 min at 30°C and the supernatant was removed. Pellet was suspended in 1.6x starting volume of ice cold BRB80 buffer to depolymerize labile microtubules. An additional 1 mM GMPCPP was added and microtubules were polymerized at 37°C for 30 min and pelleted again. The pellet was suspended in 1.6x starting volume warm BRB80. The reaction was then gently pipetted 5-10 times to shear the microtubules, aliquoted into 1.5 µL volumes, snap frozen in liquid nitrogen and stored at -80°C. Imaging chambers were assembled using 22x22 mm and 18x18 mm coverslips. Coverslips were cleaned by plasma cleaner (Harrick Plasma; Ithaca, NY) and silanized as previously described (Gell et al., 2010) and stored in desiccators at room temperature until used. Coverslips were mounted in a custom fabricated stage insert and separated by strips of parafilm. GMPCPP-stabilized microtubule seeds were affixed to coverslips using anti-rhodamine antibodies (Fisher Scientific, Cat# A-6397; diluted 1:100 in BRB80). Chambers were flushed with 1% Pluronic-F127 in BRB80 and equilibrated with an oxygen scavenging buffer (40 mM glucose, 1 mM Trolox, 64 nM Catalase, 250 nM Glucose Oxidase, 10 mg/ml Casein) prior to flowing reactions into the chambers. Purified GST-Eif4a3 (500nM) or purified GST (500nM) was added to reactions containing 15µM porcine brain tubulin labeled with 13% Hylite-488 tubulin (Cytoskeleton, Inc.), 1 mM MgCl2, 1 mM GTP, oxygen scavenging buffer in BRB80, and flowed into equilibrated chambers. Chambers were sealed with VALAP (1:1:1 Vaseline, Lanolin, Paraffin wax) and warmed to 37°C using an ASI400 Air Stream Stage Incubator (Nevtek; Williamsville, VA) for 10 minutes before imaging. Objective temperature was verified using an infrared thermometer. Images were collected on a Nikon Ti-E microscope equipped with a 1.49 NA 100× CFI160 Apochromat objective, TIRF illuminator, OBIS 488-nm and Sapphire 561-nm lasers (Coherent; Santa Clara, CA), and an ORCA-Flash 4.0 LT sCMOS camera (Hammamatsu Photonics; Japan), using NIS Elements software (Nikon; Minato, Tokyo, Japan). Images were acquired at 3 second intervals and kymographs were generated from each microtubule using a custom-made MATLAB program (Fees and Moore, 2018). Polymerization rates were calculated by measuring the changes in microtubule length and time from the beginning to the end (catastrophe) of individual polymerization events from the kymographs. Length at catastrophe was determined as the distance from the rhodamine-labeled seed to the microtubule plus at the transition from polymerization to depolymerization.

### Molecular Modeling

Schrodinger Biologics Suite 2018-3 (Schrodinger LLC, New York) was used to facilitate protein structure preparation and protein-protein docking. The structure of EIF4A3 bound to CWC22 (PDB 4C9B) was obtained from the PDB. In place of an unbound CWC22 structure, which is not available in the PDB, CWC22 was extracted from the low resolution structure of the activated human minor spliceosome (PDB 7DVQ); due to CWC22 utilising an alternative interface to binding the remaining spliceosome components to what it utilises for binding EIF4A3, as well as the lack of side chain atoms due to the low resolution of the structure (requiring these to be predicted prior to docking), this structure was deemed a suitable substitute for an unbound CWC22 structure for docking validation purposes (Bai et al., 2021). The highest resolution structure of TUBB3 available was selected from the PDB (PDB 6S8L) (La Sala et al., 2019); this structure features TUBB3 (with GDP-Mg) bound to TUBA1B (with GTP-Mg), a designed ankyrin repeat protein (DARPIN), and plinabulin. The Protein Preparation Wizard was used to prepare protein structures, with missing side-chains and loops added by Prime (Jacobson et al., 2004), ionization states of small molecules set by Epik (Shelley et al., 2007), and all water molecules deleted. For docking, the DARPIN and plinabulin were removed from PDB 6S8L, and CWC22 was removed from PDB 4C9B.

ZDOCK 3.0.2 (Pierce et al., 2011) and the pyDockRST module of pyDock3 (Chelliah et al., 2006) were used for docking and pose filtering respectively, adapting our earlier published procedures (Agostino et al., 2016). For docking purposes, EIF4A3 was defined as the receptor (fixed in position) and, in their respective docking runs, CWC22 and TUBB3-TUBA1B defined as ligands (mobile). Coarse sampling in ZDOCK was utilised, with all 3600 poses saved. EIF4A3 residues within 4.0 Å of CWC22 in PDB 4C9B were utilised as restraints within pyDockRST to facilitate pose filtering, with sequential pyDockRST runs performed to filter first for poses contacting N-terminal helicase domain residues known to contact CWC22, followed by filtering for poses contacting C-terminal helicase domain residues known to contact CWC22. Poses passing both rounds of filtering were then refined, by subjecting residues within 6.0 Å of the docked interface to Prime Side Chain Prediction followed by Prime Minimization, with the refinement process automated using a KNIME workflow (Agostino et al., 2017). Root-mean-squared deviation (RMSD) calculations on the docked CWC22 poses compared to CWC22 in the EIF4A3-CWC22 complex were performed using the Superposition tool within Maestro. The top-ranked pose following application of the docking and filtering procedure to predict the structure of the CWC22-EIF4A3 complex closely reproduced the crystal structure of this complex (Cα RMSD = 2.2 Å), suggesting the procedure’s utility for predicting the binding of proteins not induced to fit EIF4A3, as is the case for predicting tubulin binding to EIF4A3.

Following refinement, the selected top-ranked poses were subject to molecular dynamics (MD) simulations using GROMACS 2020.3 (Kohnke et al., 2020) and molecular mechanics-generalized Born/surface area (MM-GB/SA) calculations using the *MMPBSA.py* tool (Miller et al., 2012) within AmberTools 14, adapting previously published procedures (Hemming et al., 2019). Systems were parameterized for simulation in *tleap* using the *ff14SB* force field for proteins (Maier et al., 2015) and previously published AMBER parameters for nucleotide polyphosphates (Meagher et al., 2003). The resulting topologies were converted to GROMACS format using *acpype* (Sousa da Silva and Vranken, 2012)and subsequent system preparation performed in GROMACS 2020.3. Complexes were centred in a dodecahedral box with a minimum 10 Å distance to the box edge, solvated in TIP3P water, and sodium chloride added to a concentration of 0.15 M, with additional sodium ions added to charge-neutralize the system.

Following steepest descent energy minimisation of the simulation boxes, systems were briefly equilibrated (0.1ns each) in the NVT and NPT ensembles, with position restraints (1000 kJ/(mol nm^2^)) employed on all heavy atoms. A target temperature of 310K was achieved using the modified Berendsen thermostat, and a target pressure of 1 atm was achieved using the Parrinello-Rahman barostat. Short-range van der Waals and electrostatic interactions were computed to 1.2 nm, and long-range electrostatics were computed using the particle mesh Ewald approach. Constraints on bonds to hydrogens were employed using the LINCS method. Up to five simulations 50ns in length were performed for each pose, saving frames every 10ps, with three simulations selected for MM-GB/SA calculations on the basis of stabilizing in atomic coordinates within the final 10ns of the simulations. Frames from the final 10ns of the simulations were subject to MM-GB/SA calculations. The Onufriev-Bashford-Case GB model II (*igb* = 5) was employed to calculate the polar desolvation contribution (Onufriev et al., 2004), while the LCPO method was employed to calculate the nonpolar desolvation contribution. Binding energies are reported as the average energies from calculations on the three simulations.

A representative structure of the predicted tubulin-EIF4A3 complex was identified by combining the final 10ns of the three trajectories derived from the pose with the lowest binding energy by MM-GB/SA, clustering the resulting trajectory to 0.2 nm using the GROMOS algorithm (Daura et al., 1999), and selecting the middle structure of the largest cluster. Further, the MM-GB/SA binding energies derived from the trajectories of this pose were per-residue decomposed (*idecomp* = 1) to identify residues affording major contributions to the binding energy.

### Quantification and statistical analyses

All quantifications were performed blindly. Experiments were performed at least in biological triplicates unless stated otherwise. Statistical analysis was performed in GraphPad Prism software (version 8 and 9). Values are represented as mean + standard deviation (SD). The level of statistically significance was defined as P<0.05 and P values, statistical tests and sample size (n) for each experiment are informed in figure legends and Table S1. * means p≤0.05; ** means p≤0.01; *** means p≤0.001; **** means p≤0.0001

---

**Figure S1.**
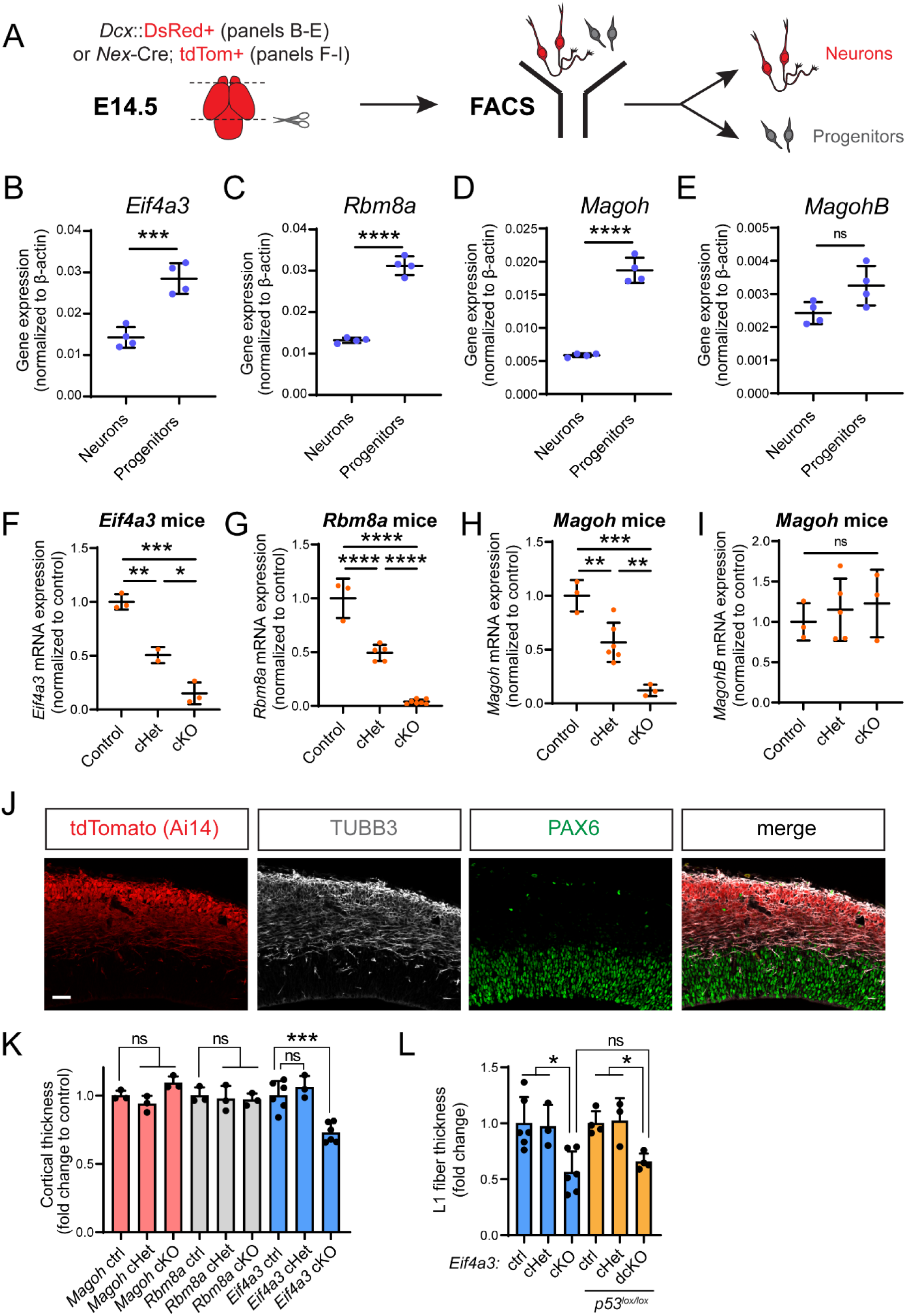
Core EJC components are expressed in cortical excitatory neurons and depleted in respective *Nex*-Cre mutants, related to Figure 1. (A), Cartoon representing Fluorescence Cell Activated Sorting (FACS) of either E14.5 DsRed positive and negative cells (from *Dcx*::DsRed mice) or E14.5 tdTomato positive cells (from *Nex*-Cre mice crossed with *Eif4a3* floxed, *Rbm8a* floxed or *Magoh* floxed mice), used to validate expression of the core EJC components. (B-E) RT-qPCR analysis of *Eif4a3* (B), *Rbm8a* (C), *Magoh* (D), *Magoh B* (F), mRNA levels in neurons and neural progenitors (DsRed positive and negative sorted cells, respectively, as described in panel A). ***p=0.0007 for Eif4a3 (B); ****p<0.0001 for Rbm8a (C) and Magoh (E), unpaired t-test (two tailed), n=4 embryos (1 litter). (F-I) RT-qPCR analysis of *Eif4a3* (F), *Rbm8a* (G), *Magoh* (H), *Magoh B* (I), mRNA levels in tdTomato positive cells sorted from indicated control, cHet or cKO EJC mutants E14.5 cortices. Each dot represents an independent embryo. (F) *p=0.0141; **p=0.0034; ***p=0.0002, one-way ANOVA (Tukey’s multiple comparison test), n=3 control embryos (2 litters), n=2 *Eif4a3* cHet embryos (1 litter) and n=3 *Eif4a3* cKO embryos (1 litter). (G) ****p<0.0001, one-way ANOVA (Tukey’s multiple comparison test), n=3 control embryos (2 litters), n=5 *Rbm8a* cHet embryos (3 litters) and n=7 *Rbm8a* cKO embryos (3 litters). (H) **p=0.008 (ctr vs. *Magoh* cHet); **p=0.0068 (*Magoh* cHet vs. *Magoh* cKO); ***p=0.0002 (ctrl vs. *Magoh* cKO), one-way ANOVA (Tukey’s multiple comparison test), n=3 control embryos (2 litters), n=6 *Magoh* cHet embryos (2 litters) and n=3 *Magoh* cKO embryos (3 litters). (I) ns=not significant. (J) Representative images of coronal section of E14.5 mouse brain from *Nex*-Cre; *Eif4a3^lox/+^*; *Ai14^lox/+^* mouse, showing tdTomato expression and stained with the neuronal marker TUBB3 (white) and the nuclear progenitor marker PAX6 (green). (K) Fold change of cortical thickness in E17.5 *Magoh*, *Rbm8a* and *Eif4a3* cHet and cKO relative to control (*Magoh*^lox/lox^, *Rbm8a*^lox/lox^ and *Eif4a3*^lox/lox^, respectively). ns=not significant; ***p=0.0004, one-way ANOVA (Dunnett’s multiple comparison test), n=3-6 embryos (from 2-3 litters). (L) Fold change of L1+ cortical axonal thickness in E17.5 *Eif4a3* dcHet and dcKO relative to control *Eif4a3*^lox/lox^ (Figure 1D) and *Eif4a3*, *p53* dcHet and dcKO relative to control *Eif4a3*^lox/lox^, *p53*^lox/lox^ (Figure 1D). *p<0.05 (see table S1 for the specific p values in each case), one-way ANOVA (Tukey’s multiple comparison test), n=4 embryos (4 litters). All graphs represent mean + S.D. Scale bars: 40 μm (J).

**Figure S2.**
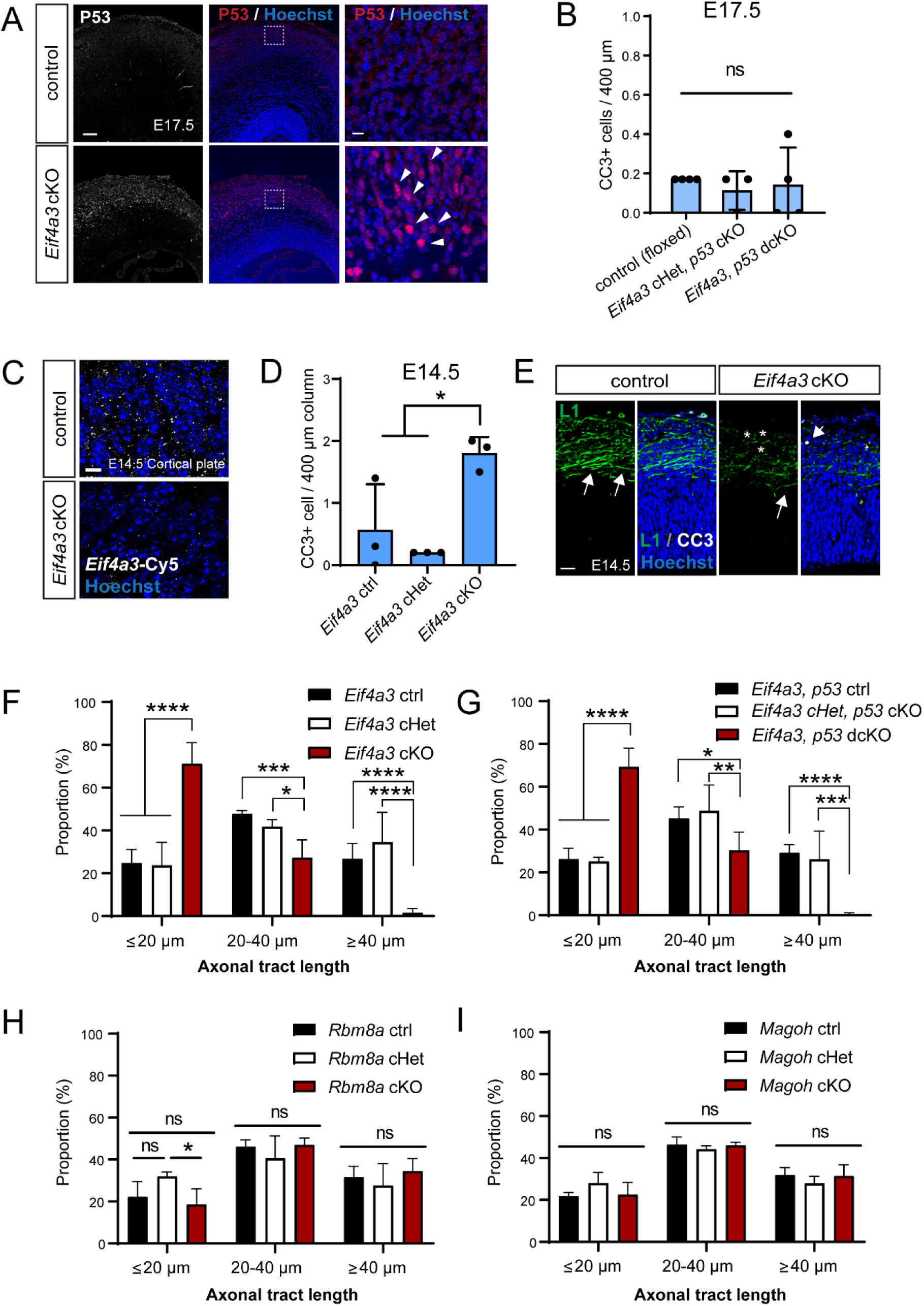
Analysis of axonal tract formation and apoptosis in EJC *Nex*-Cre mutants and *Nex*-Cre *Eif4a3, p53* double compound mutants, related to Figure 1. (A) Representative images of coronal section of E17.5 mouse brain from control (*Eif4a3*^lox/lox^) and *Eif4a3* cKO, stained with P53 (white, red) and Hoechst (blue). White boxes indicate higher magnification on the left. Arrowheads indicate examples of P53 positive nuclei. (B) Number of apoptotic cells (CC3+) per 400 μm cortical column in E17.5 *control (Eif4a3* ^lox/lox^*, p53* ^lox/lox^), *Eif4a3, p53* dcHet (*Nex*-Cre; *Eif4a3* ^lox/+^*, p53* ^lox/+^) and *Eif4a3, p53* dcKO (*Nex*-Cre; *Eif4a3* ^lox/lox^*, p53* ^lox/lox^). ns=not significant, one-way ANOVA (Tukey’s multiple comparison test), n=3-4 embryos (from 3-4 litters). (C) Representative images of *Eif4a3*-Cy5 smFISH (white dots) in E14.5 cortical plate from control (*Eif4a3*^lox/lox^) and *Eif4a3* cKO, co-stained with Hoechst (blue). (D) Number of apoptotic cells (CC3+) per 400 μm cortical column in E14.5 *control* (*Eif4a3* ^lox/lox^) *Eif4a3* cHet and *Eif4a3* cK *p=0.0359 (ctr vs. *Eif4a3* cKO); *p=0.0116 (*Eif4a3* cHet vs. *Eif4a3* cKO), one-way ANOVA (Tukey’s multiple comparison test), n=3 embryos (from 2-3 litters). (E) Representative images of coronal section of E14.5 mouse cortex from control (*Eif4a3*^lox/lox^) and *Eif4a3* cKO, stained with L1- NCAM (green), CC3 (white) and Hoechst. Groups of long and short axons are indicated with arrows and asterisks, respectively. Arrowhead shows an apoptotic nucleus. (F-I) Quantification of the proportion of axonal fibers at E14.5 categorized according to length (≤20 μm, 20-40 μm or ≥60 μm) in control, *Eif4a3* cHet and *Eif4a3* cKO (F); control, *Eif4a3, p53* dcHet and *Eif4a3, p53* dcKO (G); control, *Rbm8a* cHet and *Rbm8a* cKO (H); control, *Magoh* cHet and *Magoh* cKO (I). (F) *p=0.0247; ***p=.0006, ****p≤0.0001, one-way ANOVA (Tukey’s multiple comparison test), n=4-5 embryos (from 4 litters); (G) *p=0.0293; **p=0.0063; ***p=0.0002; ****p≤0.0001, one-way ANOVA (Tukey’s multiple comparison test), n=4 embryos (from 2-3 litters); (H) *p=0.0275, one-way ANOVA (Tukey’s multiple comparison test), n=4 embryos (2 litters); (I) ns=not significant, one-way ANOVA (Tukey’s multiple comparison test), n=2-4 embryos (2 litters) . All graphs represent mean + S.D. Scale bars: 100 (A-left) and 10 μm (A-right, C), 50 μm (E).

**Figure S3.**
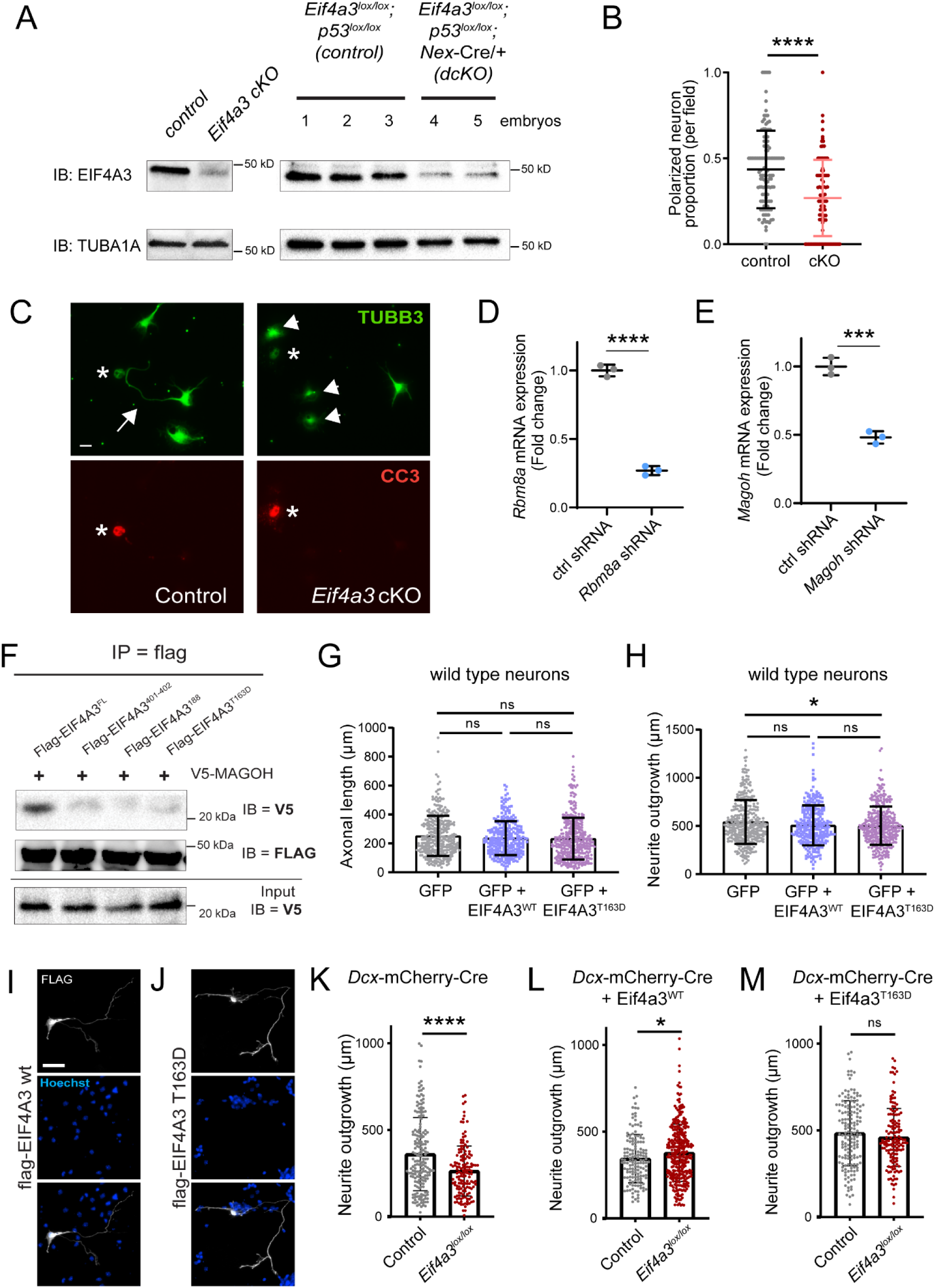
EIF4A3 is required for neuronal morphological maturation, independent of the EJC and its RNA-binding, related to Figure 2. (A) *Left:* Representative Immunoblot from DIV2 neuronal cultures from one control (*Eif4a3*^lox/lox^) and one *Eif4a3* cKO, probed for EIF4A3 and α- TUBULIN. *Right:* Representative Immunoblot from DIV2 neuronal cultures from three control (*Eif4a3*^lox/lox^, *p53*^lox/lox^) and two *Eif4a3, p53* double compound cKO (dcKO), probed for EIF4A3 and α-TUBULIN. (B) Quantification of the proportion of polarized neurons per field of low-confluency neuronal cultures from control and *Eif4a3* cKO littermates as in Figure. 2A. ****p<0.0001, unpaired t-test (two tailed), n=120 control fields (4 embryos; 3 litters) and n=90 *Eif4a3* cKO fields (3 embryos; 3 litters). (C) Representative images of neuronal cultures from E15.5 control and *Eif4a3* cKO cortices, fixed at day in vitro 2 (DIV2) and stained against CC3 (red) and TUBB3 (green). Arrow, polarized neuron. Arrowheads, unpolarized neurons. Asterisks denote infrequent apoptotic nuclei (CC3 positive), inherent to cell culture, which were not used for quantifications. (D) RT-qPCR analysis of *Rbm8a* mRNA levels in sorted N2a cells co-transfected with GFP and either a control shRNA or an *Rbm8a* shRNA. Each dot represents an independent transfected well. ****p<0.0001, unpaired t-test (two tailed), n=3 transfected wells + FAC-sorted cells. (E) RT-qPCR analysis of *Magoh mRNA* levels in sorted N2a cells co-transfected with GFP and either a control shRNA or an *Magoh* shRNA. Each dot represents an independent transfected well. ***p=0.0003, unpaired t-test (two tailed), n=3 transfected wells + FAC-sorted cells. (F) Co-immunoprecipitation between different FLAG tagged EIF4A3 proteins and V5-MAGOH co-transfected in N2a cells. Re-probing of the same blot with FLAG antibody is shown in the middle and input blot showing V5 expression is shown below. EIF4A3^401-402^ and EIF4A3^188^ mutants were included as positive controls, which also perturb EJC binding (Gehring et al., 2009). (G) (H) Axonal length (G) and neurite outgrowth (H) quantification from DIV3 neuronal cultures generated from wild type embryos *in utero* electroporated with CAG-*Gfp* and either CAG-3xFLAG-*Eif4a3*^WT^ or CAG-3xFLAG-*Eif4a3*^T163D^. ns=not significant, *p=0.0411, one-way ANOVA (Tukey’s multiple comparison test), n=335 neurons for GFP condition (from 7 embryos and 3 litters), n=283 neurons for GFP + EIF4A3^WT^ condition (6 embryos, 3 litters), n=365 neurons for GFP + EIF4A3^T163D^ condition (7 embryos, 3 litters). (I,J) Images depicting expression of Flag-EIF4A3 (I) and FLAG EIF4A3^T163D^ (J) in neurons, detected with anti-FLAG antibody. (K-M) Neurite outgrowth quantification from DIV3 neuronal cultures of control (*Eif4a3*^+/+^) or *Eif4a3*^lox/lox^ littermate embryos *in utero* electroporated with (K) CAG-GFP and *Dcx*-mCherry-Cre, (l) CAG-GFP, *Dcx*-mCherry-Cre and CAG-3x-flag- EIF4A3, and (M) CAG-GFP, *Dcx*-mCherry-Cre and CAG-3x-flag- EIF4A3^T163D^ as in Figure 2G-I. (K) ****p<0.0001. (L) *p=0.0378. (M) ns=not significant. Unpaired t-test (two tailed), n=143-310 neurons (2-5 embryos; 2-3 litters). All graphs represent mean + S.D. Scale bar: 10 μm (C), 50 μm. (I,J).

**Figure S4.**
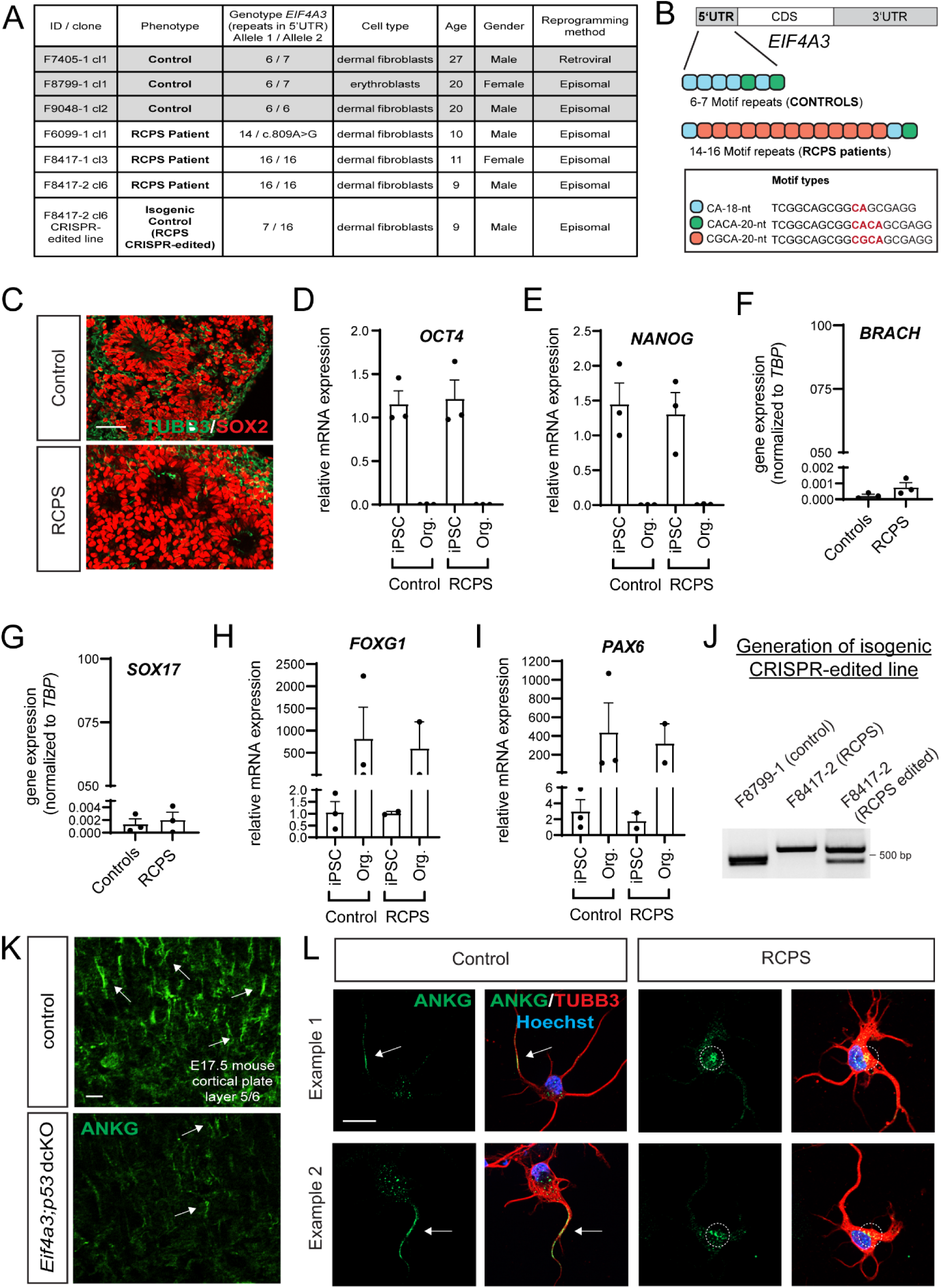
iPSC information and cortical organoid generation quality control, related to Figure 3. **(A)** Table with information of the induced pluripotent stem cells (iPSCs) used in this study. Adapted from Miller *et al*. 2017. **(B)** Cartoon representing 5’ UTR region of EIF4A3 in control and RCPS patients which present non-coding pathological expansion. Unaffected individuals typically have 6-9 repeats composed mostly by CA-18nt- and CACA-20nt-long motifs within the EIF4A3 5’UTR. In contrast, of 42 affected RCPS individuals identified so far, most are homozygous for 14-16 repeats and the majority of them are the disease-associated CGCA-20nt motif, resulting in hypomorphic *EIF4A3* expression. **(C)** Images of cortical organoids at day 35 and stained for TUBB3 (green) and SOX2 (red). **(D-I)** RT-qPCR analysis of mRNA levels for *OCT4* (**D**), *NANOG* (**E**), *BRACH* (**F**), *SOX17* (**g**), *FOXG1* (**H**), and *PAX6* (**I**), in control and RCPS iPSC lines and cortical organoids. Each dot represents a different cell line whose value was obtained by averaging the data coming from 3-9 organoids from two independent differentiations. All organoids were from D25. **(J)** Gel representing generation of isogenic line. Note two bands are now detected in the CRISPR edited line reflecting the RCPS allele with repeats and a corrected WT allele. **(K)** Representative images of coronal section of E17.5 mouse cortical plate from control (*Eif4a3*^lox/lox^; *p53*^lox/lox^) and *Eif4a3* dcKO (*Nex*-Cre; *Eif4a3*^lox/lox^; *p53*^lox/lox^), stained with ANKG. Arrows indicate AIS observed in longitudinal plane. **(L)** Additional representative images from DIV7 neurons of control and RCPS patient cultures generated from dissociated cortical D71-73 organoids, stained with ANKG, TUBB3 and Hoechst, as in Figure 3. Scale bar: 10 um (J), 20 μm (L).

**Figure S5.**
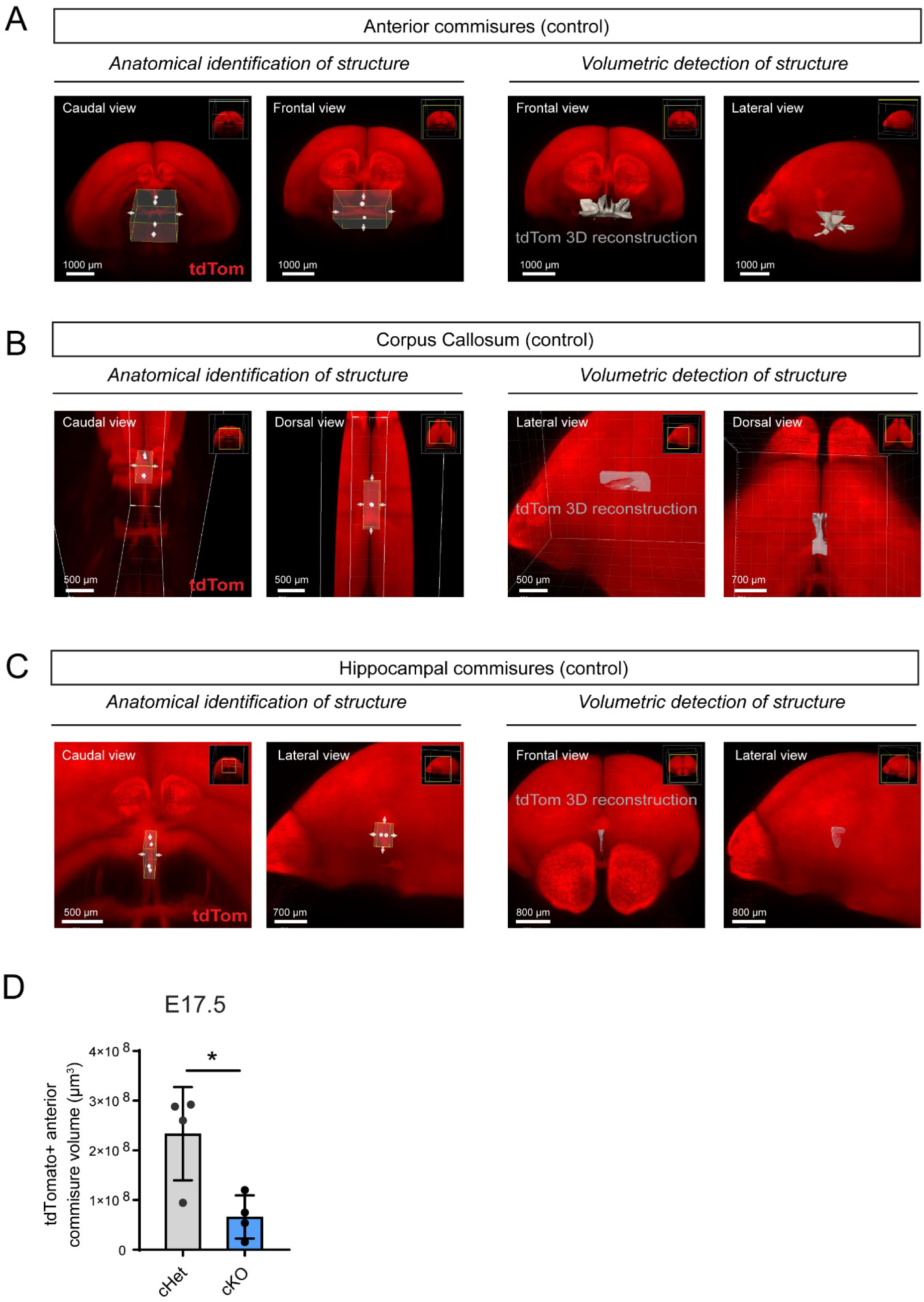
Anatomical identification of the three main intracortical axonal tracts in 3D reconstructed brains cleared by CUBIC, related to Figure 4. (A-C) Representative 3D reconstructions of tdTomato positive E17.5 cleared brains (by CUBIC) from control (*Eif4a3* cHet) to identify and quantify the volume of the anterior commissures (A), corpus callosum (B), and the hippocampal commissures (C). Yellow boxes with white arrows are the region of interest that were defined to detect each structure. Grey displays indicate the volumetric detection of the region of interest. Brains were positioned in caudal, frontal, lateral or dorsal view to identify each structure. (D) Anterior commissures volume in E17.5 *Eif4a3* cHet (control) and *Eif4a3* cKO brains. *p=0.0177, unpaired t-test (two tailed), n=4 control and *Eif4a3* cKO brains (2 litters). Scale bars are embedded on each image. All graphs represent mean + S.D.

**Figure S6.**
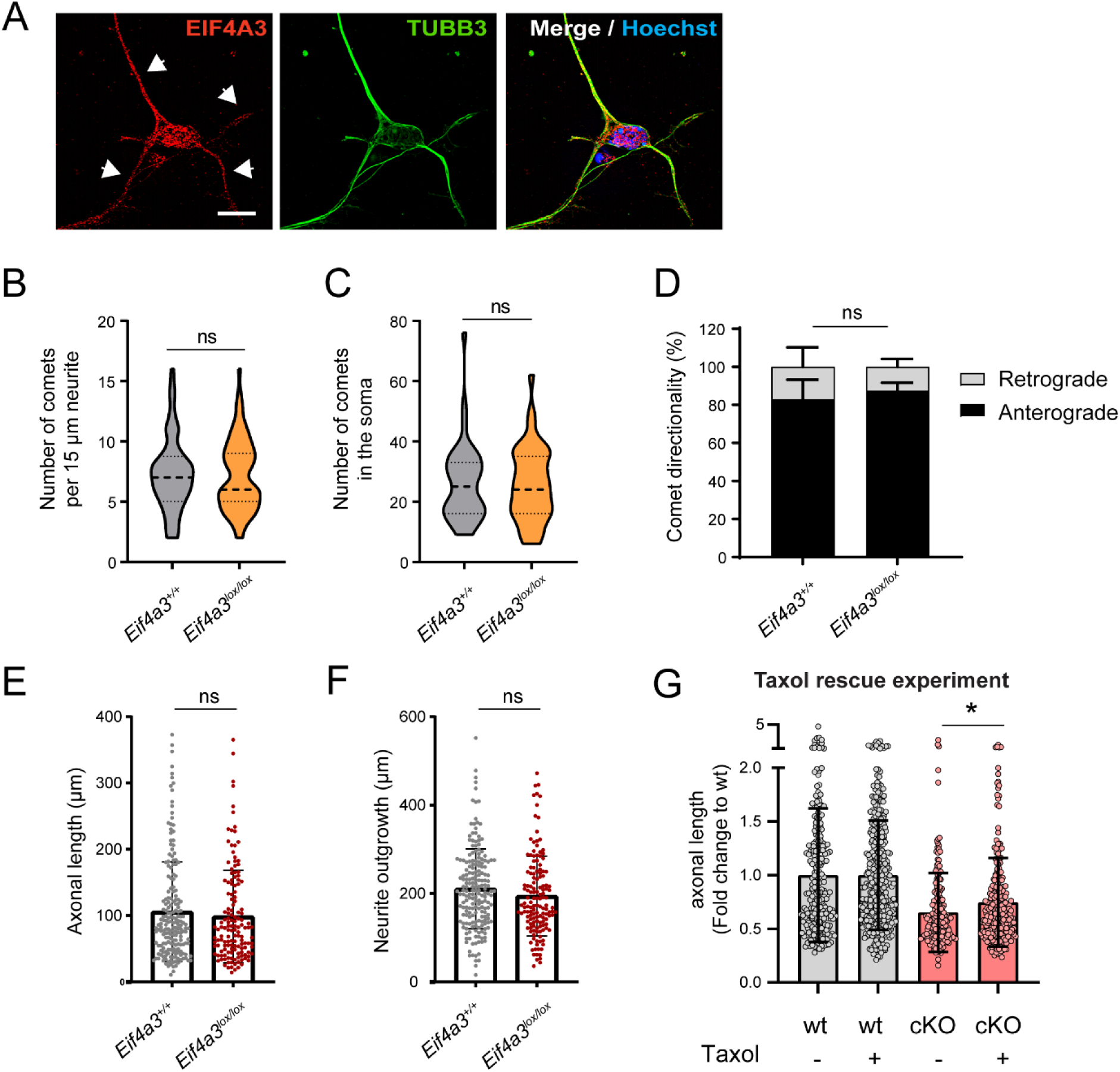
EIF4A3 is localized in neuronal projections and number of microtubule comets is not affected in *Eif4a3* cKO neurons, related to Figure 5. (A) Representative SIM super-resolution images of a cortical neuron from wild type E15.5 neuronal cultures, fixed at DIV2 and stained against either EIF4A3 (red, independent antibody from Figures 5A and 5C), TUBB3 (green) and Hoechst (blue). Arrows denote neuronal projections. (B,C) Number of comets per 15 μm of neurites (B) and number of comets in the soma (C), at time=0 of the live imaging, of three independent neuronal cultures from electroporated control (*Eif4a3*^+/+^) or *Eif4a3*^lox/lox^ littermates generated as in Figure 5D. ns=not significant, unpaired t-test (two tailed), (B) n=80 control neurites (from 41 electroporated neurons, 4 embryos, 3 litters) and n=81 *Eif4a3* lox/lox neurites (from 41 electroporated neurons, 4 embryos, 3 litters). (C) n=41 control and *Eif4a3* lox/lox electroporated neurons (4 embryos, 3 litters). (D) Comet directionality of three independent neuronal cultures from electroporated control (*Eif4a3*^+/+^) or *Eif4a3*^lox/lox^ littermates generated as in Figure 5D. ns=not significant, two-way ANOVA (Šídák’s multiple comparison test), n=15 control electroporated neurons (1307 traced comets from 2-6 neurites per neuron, 3 embryos and 2 litters) and n=20 *Eif4a3* lox/lox electroporated neurons (2049 traced comets from 2-6 neurites per neuron, 4 embryos and 3 litters). (E,F) Axonal length (E) and neurite outgrowth (F) quantification from neuronal cultures of control (*Eif4a3*^+/+^) or *Eif4a3*^lox/lox^ littermate embryos *in utero* electroporated with CAG-GFP and *Dcx*-mCherry-Cre as described in Figure 2 and fixed at DIV2 (same DIV when live imaging was performed). ns=not significant, unpaired t-test (two tailed), n=192 electroporated control neurons (5 embryos, 3 litters) and n=143 electroporated *Eif4a3* lox/lox neurons (4 embryos, 3 litters). (G) Axonal length quantification from neuronal cultures of electroporated control (grey) or *Eif4a3* mutant (red) and treated with Taxol. p=0.0104, unpaired t-test (two tailed), n=243 neurons for control no Taxol (from 3 embryos, 1 litter), n=210 neurons for control with Taxol (3 embryos, 1 litter), n=365 neurons for *Eif4a3* lox/lox no Taxol (3 embryos, 1 litter) and n=238 neurons for *Eif4a3* lox/lox with Taxol (3 embryos, 1 litter). All graph bars represent mean + S.D.; truncated violin plots show median (thicker dashed-line) ± quartiles (thinner dashed-lines) Scale bar: 10 μm (A).

**Figure S7.**
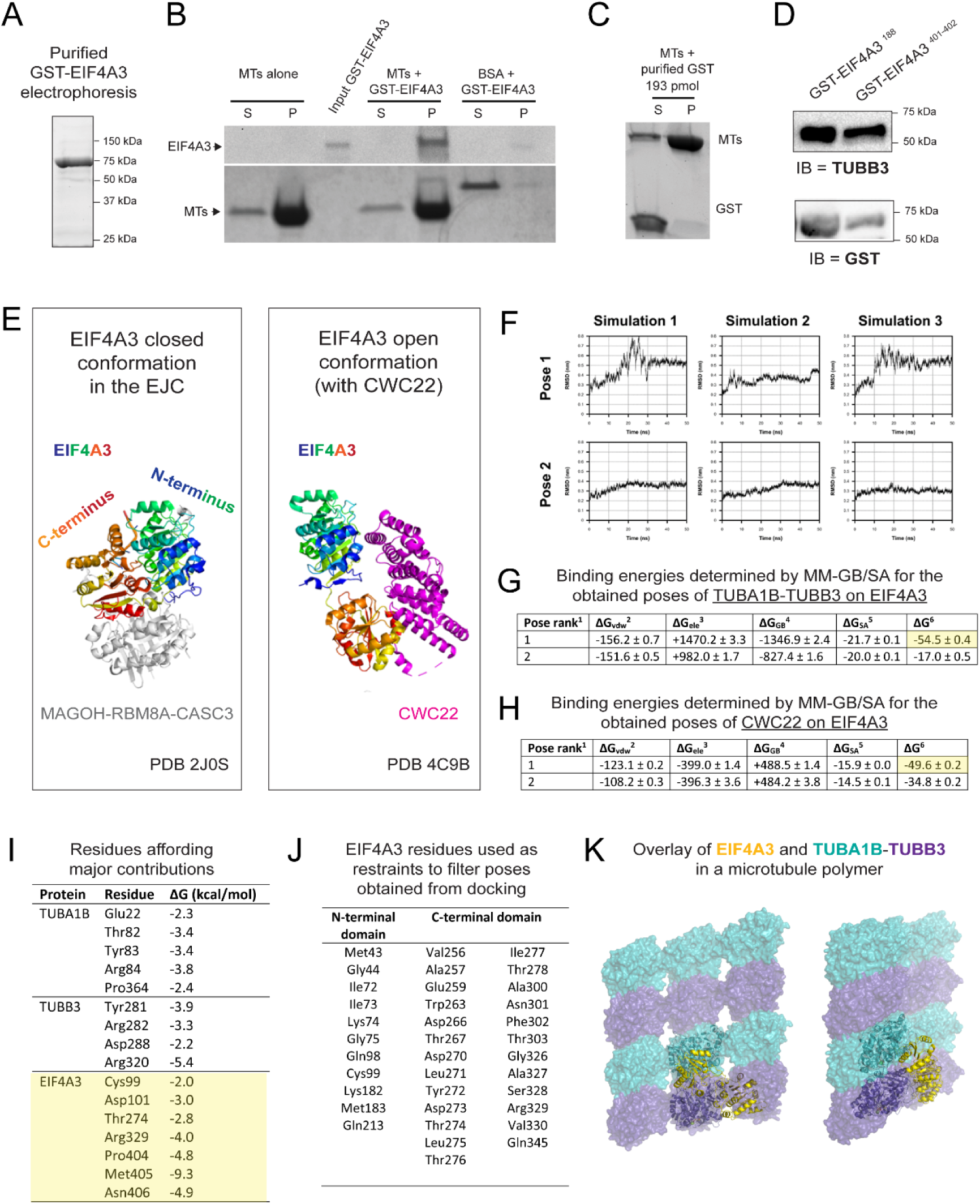
EIF4A3 directly binds to microtubules in a different conformation and with different residues than the EJC, related to Figure 6 and 7. (A) Stain-free gel image depicting purified GST- EIF4A3. (B) Representative western blot depicts co-sedimentation of purified GST- EIF4A3 or GST alone with Taxol-polymerized microtubules, probed for EIF4A3 and α-TUBULIN. Each lane was loaded with the entire pellet (P), 1/3 of supernatant, and input was 1/10 of total lysate from reaction. Absence of EIF4A3 in the supernatant is due to the diluted sample also containing sucrose cushion (S). (C) Co-sedimentation of purified GST with polymerized microtubules (total protein gel). (D) Representative western blot of a GST pulled down experiment (as in Figure 6F), where microtubules were incubated with either mutant EIF4A3^188^ and EIF4A3^401-402^. Blot against TUBB3 is shown on the left. Re-probing of GST is shown on the right. (N=2 experiments). (E) Representative structures of EIF4A3 from the PDB. Left, EIF4A3 bound to AMPPNP-Mg, an RNA fragment, CASC3, MAGOH and RBM8A (PDB 2J0S). Right, EIF4A3 bound to CWC22 (PDB 4C9B). Structures are aligned using the N-terminal domain of EIF4A3 (which is oriented identical in both cases). with the approximate position of the C-terminal domain in each structure indicated by the label. Legend: blue-to-red rainbow – N-to-C-terminus of EIF4A3 (with N-terminal domain colored blue-to-green, and C-terminal domain colored yellow-to-red); white cartoon/transparent surface – CASC3, MAGOH and RBM8A; pink cartoon/transparent surface – CWC22. (F) RMSDs over time for molecular dynamics simulations of tubulin docked to EIF4A3. (G, H) Binding energies (in kcal/mol) determined by MM-GB/SA for the obtained poses of TUBB3-TUBA1B (PDB 6S8L) on EIF4A3 (PDB 4C9B) (G) or CWC22 (PDB 7DVQ) on EIF4A3 (PDB 4C9B) (H). ^1^As determined by ZDOCK, following contact-based filtering of poses. ^2^Molecular mechanics van der Waals term. ^3^Molecular mechanics electrostatics terms. ^4^Polar desolvation from generalized Born model. ^5^Nonpolar desolvation from LCPO model. ^6^Binding energy, as given by the sum of the preceding component terms. (I) Residues affording major contributions (-2.0 kcal/mol or greater in magnitude) to the binding energy between TUBB3- TUBA1B and EIF4A3. (J) EIF4A3 residues used as restraints to filter poses obtained from docking. (K) Overlay of TUBB3-TUBA1B-EIF4A3 complex obtained from docking to a fragment of a representative microtubule structure (PDB 6DPU). Left, view of inner microtubule surface. Right, Side view of microtubule (rotated approximately 45 degrees with respect to left panel. Legend: Teal - alpha-tubulin; violet - beta-tubulin; yellow - EIF4A3.

## Video S1

3D reconstruction of CUBIC-cleared mouse heads from E14.5 *Nex*-Cre; *Eif4a3*^lox/+^; *Ai14*^lox/+^ (control) or *Nex*-Cre; *Eif4a3*^lox/lox^; *Ai14*^lox/+^ (cKO) littermate embryos. The main intracortical axonal tracts forming at this stage (the anterior commissures) are indicated for each genotype.

## Video S2

3D reconstruction of CUBIC-cleared mouse heads from E17.5 *Nex*-Cre; *Eif4a3*^lox/+^; *Ai14*^lox/+^ (control) or *Nex*-Cre; *Eif4a3*^lox/lox^; *Ai14*^lox/+^ (cKO) littermate embryos. The three main intracortical axonal tracts forming at this stage, namely corpus callosum, anterior commissures and hippocampal commissures, are indicated for each genotype.

## Video S3

3D reconstruction of CUBIC-cleared E17.5 mouse brains from *Eif4a3*^+/+^ (control) or *Eif4a3*^lox/lox^ littermate embryos, *in utero* electroporated (IUEd) at E14.5 with *Dcx*-Cre-GFP and lox-stop-lox-tdTomato (Cre reporter). Electroporated cortices showing sparsely labeled neurons and their axons extending to the midline (callosal projections) are indicated for each genotype.

## Video S4

Time lapse imaging of growing microtubules (GFP-MACF43) in day *in vitro* 2 (DIV2) neurons, prepared from E15.5 brain cortices from *Eif4a3*^+/+^ (control) or *Eif4a3*^lox/lox^ littermate embryos, which were *in utero* electroporated at E14.5 with GFP-MACF43 and *Dcx*-mCherry-Cre. Individual GFP+ comets moving through neuronal projections (neurites) are indicated for each genotype. Images were acquired at 0.5 sec (2 fps) and video is displayed at 5x speed (10 fps).

## Notes

### Competing Interest Statement

The authors have declared no competing interest.

